# Endogenous protein tagging in medaka using a simplified CRISPR/Cas9 knock-in approach

**DOI:** 10.1101/2021.07.29.454295

**Authors:** Ali Seleit, Alexander Aulehla, Alexandre Paix

**Affiliations:** Developmental Biology Unit, European Molecular Biology Laboratory, Heidelberg, Meyerhofstrasse 1, 69117, Heidelberg, Germany

**Keywords:** CRISPR, HDR, medaka, fusion-proteins, WGS, cloning-free, Pcna, KI

## Abstract

The CRISPR/Cas9 system has been used to generate fluorescently labelled fusion proteins by homology directed repair in a variety of species. Despite its revolutionary success, there remains an urgent need for increased simplicity and efficiency of genome editing in research organisms. Here, we establish a simplified, highly efficient and precise strategy for CRISPR/Cas9 mediated endogenous protein tagging in medaka (*Oryzias latipes*). We use a cloning-free approach that relies on PCR amplified donor fragments containing the fluorescent reporter sequences flanked by short homology arms (30-40bp), a synthetic sgRNA and streptavidin tagged Cas9. We generate six novel knock-in lines with high efficiency of F0 targeting and germline transmission. Whole Genome Sequencing (WGS) results reveal single-copy integration events only at the targeted *loci*. We provide an initial characterization of these fusion-protein lines, significantly expanding the repertoire of genetic tools available in medaka. In particular, we show that the *mScarlet-pcna* knock-in line has the potential to serve as an organismal-wide label for proliferative zones and an endogenous cell cycle reporter.

## Introduction

The advent of gene editing tools (Wang et al., 2016, Jinek et al., 2012, Cong et al., 2013) in conjunction with the expansion of sequenced genomes and engineered fluorescent proteins (Chudakov et al., 2010, Shaner et al., 2013, Bindels et al., 2017) has revolutionized the ability to generate endogenous fusion protein Knock-In (KI) lines in a growing number of organisms (Paix et al., 2015, Paix et al., 2017b, Paix et al., 2017a, Gratz et al., 2014, Kanca et al., 2019, Wierson et al., 2020, Gutierrez-Triana et al., 2018, Hisano et al., 2015, Auer and Del Bene, 2014, Yoshimi et al., 2016, Yao et al., 2017, Cong et al., 2013, Dickinson et al., 2015, Leonetti et al., 2016). These molecular markers expressed at physiological levels are central to our understanding of cellular and tissue level dynamics during embryonic development (Gibson et al., 2013). To this end researchers have utilized the *Streptococcus pyogenes* CRISPR associated protein 9 (Cas9) and a programmed associated single-guide RNA (sgRNA) to introduce a Double Strand Break (DSB) at a pre-defined genomic location (Jinek et al., 2012, Cong et al., 2013). Cell DNA repair mechanisms are triggered by the DSB and it has been shown that providing DNA repair donors with homology arms that match those of the targeted *locus* can lead to integration of the donor constructs containing fluorescent reporter sequences in the genome by the process of Homology Directed Repair (HDR) (Danner et al., 2017, Jasin and Haber, 2016, Ceccaldi et al., 2016, Hoshijima et al., 2016, Shin et al., 2014, Zu et al., 2013). Despite its success, HDR mediated precise single-copy KI efficiencies in vertebrate models can still be low and the process of generating KI lines remains cumbersome and time-consuming. Recent reports have improved the methodology by the usage of 5’ biotinylated long homology arms that prevent concatemerization of the injected dsDNA (Gutierrez-Triana et al., 2018) or by linking the repair donor to the Cas9 protein (Gu et al., 2018). In addition, repair donors with shorter homology arms in combination with *in vivo* linearization of the donor plasmid have been shown to mediate efficient Knock-Ins in Zebrafish and in mammalian cells (Wierson et al., 2020, Hisano et al., 2015, Cristea et al., 2013, Auer et al., 2014, Ota et al., 2014, Shin et al., 2014).

In this work we establish a simplified, highly efficient and precise strategy for CRISPR/Cas9 mediated endogenous protein tagging in medaka (*Oryzias latipes*). Our approach relies on the use of biotinylated PCR amplified donor fragments that contain the fluorescent reporter sequences flanked by short homology arms (30-40bp), by-passing the need for cloning or *in vivo* linearisation. We use this approach to generate and characterize a series of novel knock-in lines in medaka fish (Table 1 and Table S1). By utilizing Whole Genome Sequencing (WGS) with high coverage in conjunction with Sanger sequencing of edited *loci*, we provide strong evidence for precise single-copy integration events only at the desired *loci*. In addition to generating an endogenous ubiquitous nuclear label and novel tissue specific reporters, the knock-in lines allow us to record cellular processes, such as intracellular trafficking and stress granule formation in 4D during embryonic development, significantly expanding the genetic toolkit available in medaka. Finally, we provide proof of principle evidence that the endogenous *mScarlet-pcna* knock-in we generate serves as a *bona fide* proliferative cell label and an endogenous cell cycle reporter, with broad application potential in a vertebrate model system.

## Results

### A simplified, highly efficient strategy for CRISPR/Cas9 mediated fluorescent protein knock-ins in medaka

To simplify the process of generating fluorescent protein knock-ins in teleosts we utilized PCR amplified dsDNA donors with short homology arms (30-40bp). Biotinylated 5’ ends were used to prevent *in vivo* concatemerization (Gutierrez-Triana et al., 2018, Auer et al., 2014, Winkler et al., 1991) and a Strepdavidin tagged Cas9 (Cas9-mSA) was used to increase binding affinity to the biotinylated donor constructs (Gu et al., 2018). This approach by-passes the need for cloning, as homology arms are already included in the amplification primers, and the use of a second gRNA for *in vivo* plasmid linearisation (Hoshijima et al., 2016, Shin et al., 2014, Zu et al., 2013). The three-component mix; biotinylated PCR amplified dsDNA donors, synthetic sgRNA and Cas9-mSA mRNA (Tables S2/S3/S4/S5) was injected into 1 cell-stage medaka embryos (Figure 1 and Figure S1, for a detailed protocol see Files S1 and S2). We targeted a list of seven genes with a variety of fluorescent proteins (Figure 1, Figure S1, Table 1 and Table S4), both N and C terminus tags were attempted (a list of all genomic *loci* targeted can be found in Table S1). Targeting efficiency in F0 ranged from 11% to 59% of embryos showing mosaic expression (Table 1 and Table S1). Control injections with the *actb* sgRNA, Cas9-mSA mRNA and the donor eGFP construct without homology arms showed no evidence of eGFP positive cell clones in F0 (Table S1), while the same construct with homology arms resulted in 39% of surviving injected embryos showing mosaic expression of eGFP (Table 1 and Table S1). The germline transmission efficiency of fluorescent F0 fish ranged from 25% to 100% for the different targeted *loci* (Table 1 and Table S1). For F0s with germline transmission the range of positive F1 embryos obtained was between 6.6% and 50%. Using this method we were able to establish six stable knock-in lines. Importantly, a single injection round was sufficient to generate a knock-in line for most targeted *loci* (5/6; Table 1 and Tables S1/S6). As previously reported, the *actb-eGFP* tag was embryonic lethal (Gutierrez-Triana et al., 2018) and we could not obtain a knock-in line for that *locus*. Combined, our results provide evidence that highly efficient targeting of endogenous *loci* with large inserts (∼800bp) is possible in medaka using our simplified KI approach (Figure 1 and Figure S1). In addition to being highly efficient, our method is rapid and simple-to-implement, as it relies on PCR amplification of the linear donor construct and hence alleviates the need for any additional cloning or *in vivo* plasmid linearization (Figure S1).

**Figure 1:**
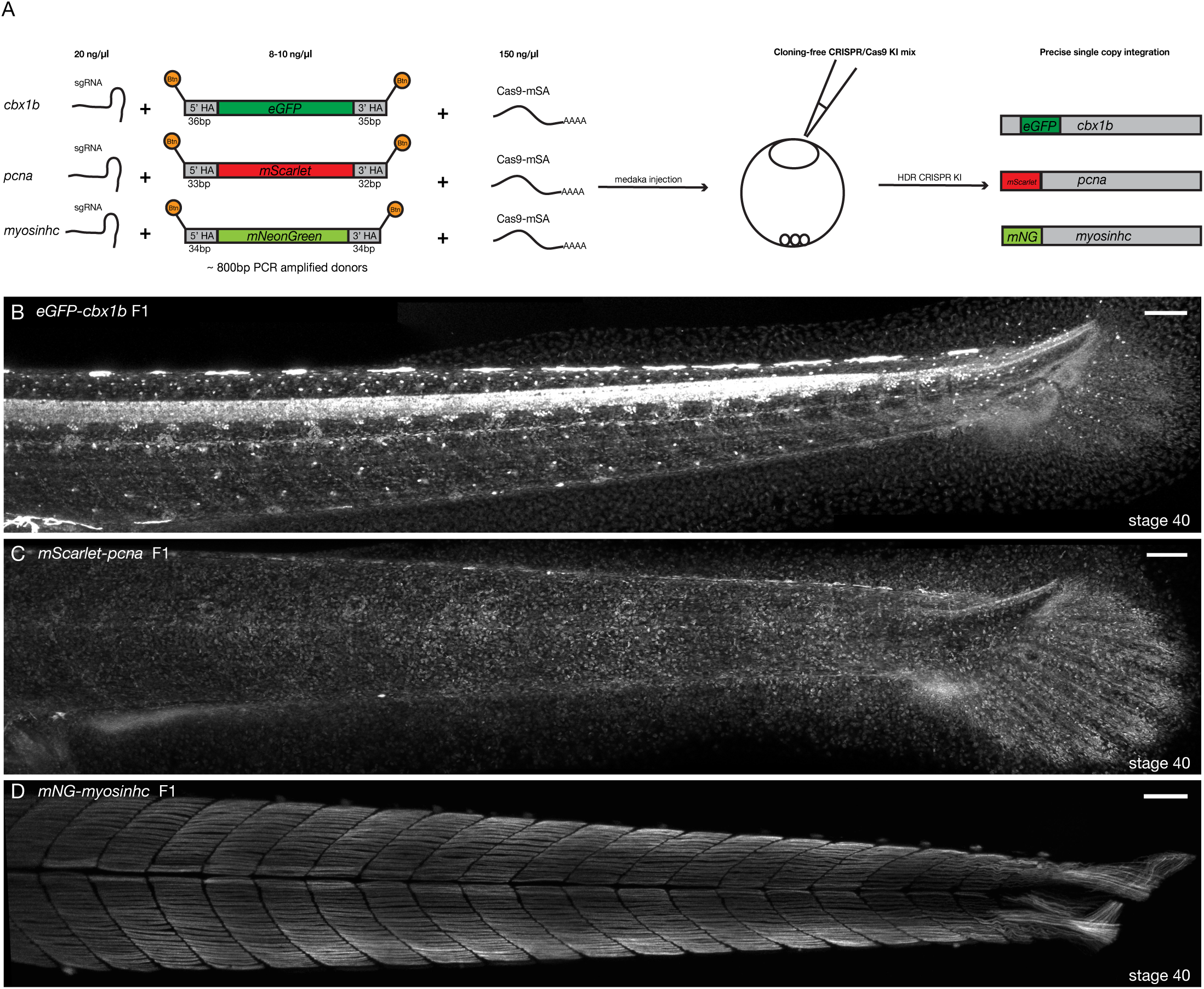
Cloning-free single copy CRISPR/Cas9 mediated KI lines in medaka. (**A**) Schematic diagram of cloning-free CRISPR knock-in strategy. The injection mix consists of three components, a sgRNA targeting the gene of interest, Cas9-mSA mRNA and the PCR amplified donor plasmid containing short homology arms on both ends (30-40bp) and the fluorescent protein of interest with no ATG and no Stop codon. Note that the 5’ ends of the PCR donor fragment are biotinylated (Btn). The mix is injected in one-cell staged medaka embryos and the injected fishes are screened for potential in-frame integrations mediated by Homology-Directed Repair (HDR). (**B**) *eGFP-cbx1b* F1 CRISPR KI line stage 40 medaka embryos. eGFP-Cbx1b labels all nuclei. n>10 embryos. Scale bar = 100µm (**C**) *mScarlet-pcna* F1 CRISPR KI line stage 40 medaka embryos. mScarlet-Pcna labels exclusively cycling cells. n>10 embryos. Scale bar = 100µm. (**D**) *mNG-myosinhc* F1 CRISPR KI line stage 40 medaka embryos. line. mNG-Myosinhc labels exclusively muscle cells. n>10 embryos. Scale bar = 100µm.

### Precise, single copy Knock-Ins of fluorescent protein reporters

We next assessed the specificity and precision of the approach. It is possible that either concatemerization of inserts or off-target integrations could occur after foreign DNA delivery and a CRISPR/Cas9 mediated DSBs (Gutierrez-Triana et al., 2018, Doench et al., 2016, Fu et al., 2013, Paix et al., 2017a, Won and Dawid, 2017, Yan et al., 2013, Hackett et al., 2007, Wierson et al., 2020, Wierson et al., 2019). To identify off-target insertions genome-wide and verify single-copy integration we performed next generation Whole Genome Sequencing (WGS) with high coverage (for details of WGS, see material and methods) on three knock-in lines (Figure 1B-D); *eGFP-cbx1b, mScarlet-pcna* and *mNeonGreen-myosinhc* . For the *eGFP-cbx1b* KI line, we could only identify paired-end *eGFP* reads anchored to the endogenous *cbx1b locus* and nowhere else in the genome (Figure S2). Likewise, in the *mScarlet-pcna* line*, mScarlet* reads only mapped to the endogenous *pcn*a *locus* (Figure S2). For the *mNeonGreen-myosinhc* line*, mNeonGreen* sequences mapped to the *myosinhc locus* (Figure S2), but paired-end analysis yielded a second, weakly supported partial insertion of *mNeonGreen* at an intronic region in the *edf1* gene. We were not able to confirm the latter insertion by subsequent PCR and hence it remains unclear whether this a false positive prediction or a mosaic insertion of very low frequency. Combined, the whole genome sequencing results therefore provide strong evidence that the method we report results in single-copy insertions only at the targeted *locus*. In addition to WGS, genotyping F1 adults followed by Sanger sequencing confirmed the generation of single-copy in-frame fusion proteins in the *eGFP-cbx1b, mScarlet-pcna and mNeonGreen-myosin-hc* lines (Figure S2). In 5/6 cases homology directed repair (HDR) resulted in precise, scarless integrations (Figure S2), while in one case we could detect a partial duplication (21 base pairs) within the 5’ homology arm, 22 base pairs upstream of the start codon (for details see Materials and Methods). Overall, the method we present here shows high precision and specificity enabling the rapid generation of endogenously tagged alleles in a vertebrate model.

### Visualisation of endogenous protein dynamics enables *in vivo* recording of cellular processes in medaka

As a proof of principle, we employed the simplified CRISPR/Cas9 strategy to generate a series of endogenous fusion protein knock-in medaka lines (Table 1 and Tables S1/S6, Figures S3/S4/S5/S6). Here, we provide an initial characterization of six of these novel knock-in lines that are made available to the community, to label cell compartments (nucleus), cell processes (cell cycle, intra-cellular trafficking, stress granule formation), cell-adhesion (adherens junctions) and specific cell types (muscle cells).

#### Ubiquitous nuclear marker

To generate a ubiquitously expressed nuclear label reporter line, we targeted the *cbx1b* (Chromobox protein homolog) *locus* with eGFP. Cbx1b is a member of the chromobox DNA binding protein family and is a known component of heterochromatin that is expressed ubiquitously (Lomberk et al., 2006, Nielsen et al., 2001, Gilmore et al., 2016). Chromobox proteins are involved in several important functions within the nucleus, such as transcription, nuclear architecture, and DNA damage response (Luijsterburg et al., 2009). We generated an *eGFP-cbx1b* KI by targeting *eGFP* to the N-terminus of the *cbx1b* coding sequence in medaka. The resulting line expresses eGFP in all nuclei of every tissue examined, and serves as the first endogenous ubiquitous nuclear label in teleosts (Figures 1B/2A, Figure S3 and Supplementary Movie 1, n>10 embryos).

**Figure 2:**
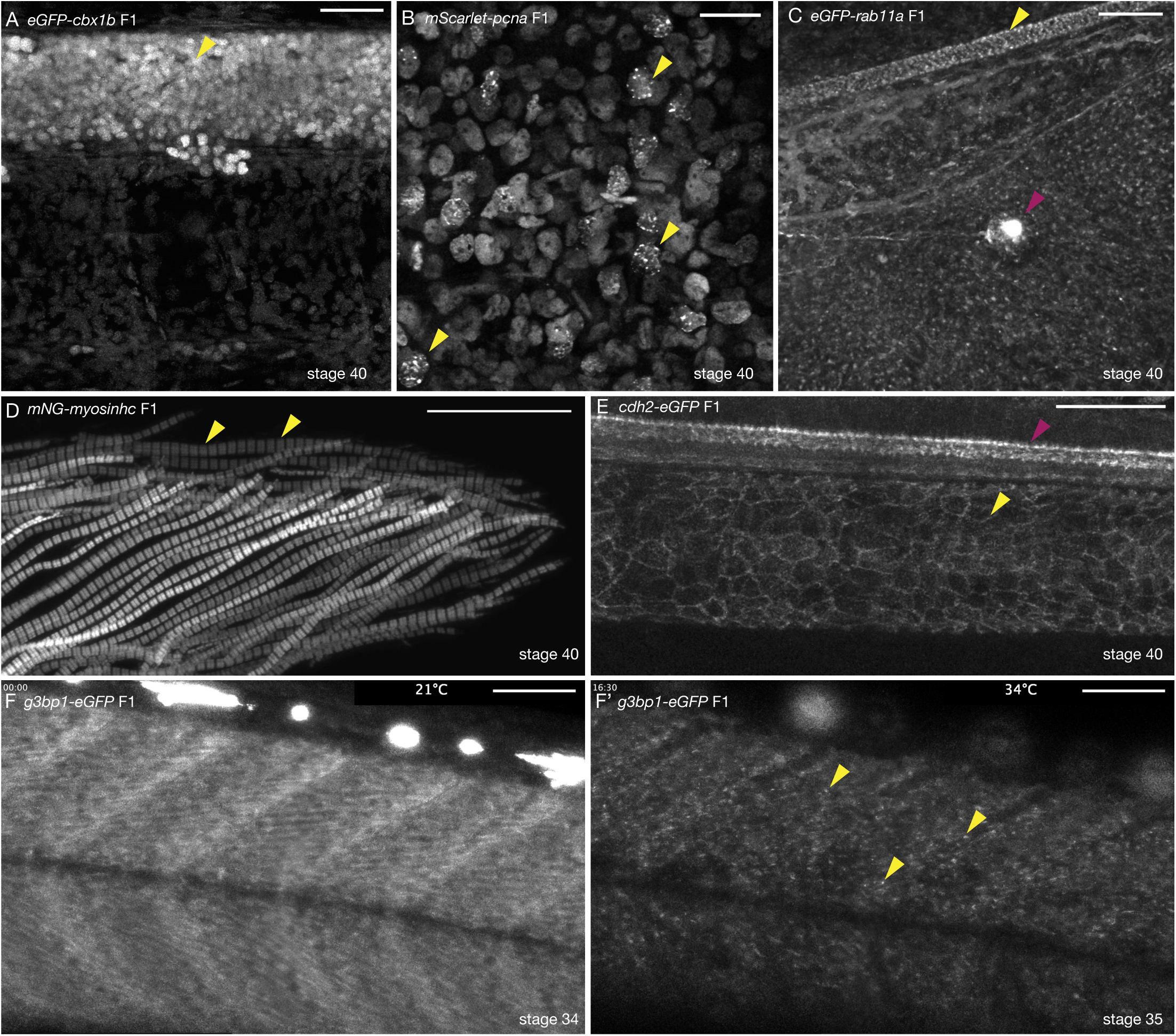
Tissue and organelle specific expression of six CRISPR/Cas9 KI lines in medaka. (**A**) *eGFP-cbx1b* F1 stage 40 medaka embryo. eGFP-Cbx1b labels all nuclei. Nuclei in the spinal cord of medaka are highlighted (Yellow arrowhead). n>10 embryos. Scale bar = 30µm. (**B**) *mScarlet-pcna* F1 stage 40 medaka embryo. mScarlet-Pcna is localized in the nuclei of cycling cells. mScarlet-Pcna is visible in skin epithelial cell nuclei located in the mid-trunk region of a medaka embryo. The localization of Pcna as speckles within the nucleus indicates cells in S phase of the cell cycle (Yellow arrowheads). n= 10 embryos. Scale bar = 20µm. (**C**) *eGFP-rab11a* F1 stage 40 medaka embryo. Expression of the membrane trafficking marker eGFP-Rab11a is evident in the caudal fin region. eGFP-Rab11a is strongly expressed in the spinal cord (Yellow arrowhead) and lateral line neuromasts (Magenta arrowhead). n=6 embryos. Scale bar = 30µm. (**D**) *mNG-myosinhc* F1 stage 40 medaka embryo. mNG-Myosinhc is expressed solely in muscle cells. Myofibrils containing chains of individual sarcomere can be seen (mNG-Myosinhc labels the Myosin A band inside each sarcomere). n>10 embryos. Scale bar = 30µm. (**E**) *cdh2-eGFP* F1 stage 40 medaka embryo. Cdh2-eGFP is localized at cell membranes in several tissues, including the spinal cord (Magenta arrowhead) and the notochord (Yellow arrowhead). n= 5 embryos. Scale bar= 50µm. (**F-F’**) *g3bp1-eGFP* F1 stage 34-35 medaka embryo. Time-lapse imaging of G3bp1-eGFP dynamics under normal and stress conditions. (F) G3bp1-eGFP localizes to the cytoplasm under physiological conditions. (F’) under stress conditions (temperature shock), G3bp1-eGFP localizes to stress granules (Yellow arrowheads). Time in hours. n= 8 embryos. Scale bar= 50µm.

#### Proliferative cell marker

With the goal of generating an endogenous cell cycle reporter, we targeted the *pcna* (Proliferating cell nuclear antigen) *locus* to generate a *mScarlet-pcna* fusion protein. Pcna is an essential protein regulator of DNA replication and integrity in eukaryotic cells (Moldovan et al., 2007, Maga and Hubscher, 2003, Mailand et al., 2013). It has been previously shown that cells that exit the cell cycle, *e.g.* post-mitotic differentiated cell types, express very low levels of Pcna (Zerjatke et al., 2017, Thacker et al., 2003, Yamaguchi et al., 1995, Buttitta et al., 2010). This has led researchers to utilize Pcna as a highly conserved marker for proliferating cells (Zerjatke et al., 2017, Barr et al., 2016, Leonhardt et al., 2000, Leung et al., 2011, Piwko et al., 2010, Alunni et al., 2010). In addition to being a specific label for cycling cells, the appearance of nuclear speckles of Pcna within the nucleus is a hallmark of cells in late S phase of the cell cycle (Zerjatke et al., 2017, Barr et al., 2016, Leonhardt et al., 2000, Leung et al., 2011, Piwko et al., 2010, Santos et al., 2015). More recently, endogenously tagged Pcna has been used in mammalian cell lines to dynamically score all the different cell cycle stages (Zerjatke et al., 2017, Held et al., 2010, Piwko et al., 2010, Santos et al., 2015). We targeted the first exon of *pcna* with *mScarlet* with high efficiency (28% mosaic expression in F0s, and 50% germline transmission) and generated the *mScarlet-pcna* KI line (Figures 1C/2B, n>10 embryos). Using stage 40 medaka embryos, we detected mScarlet-Pcna positive cells within the epidermis, specifically in supra-basal epidermal cells (Figure 2B). A subset of these cells showed nuclear speckles of mScarlet-Pcna that likely represent replication foci and are a characteristic marker for late S phase (Figure 2B, yellow arrowheads). We validate the use of this line both as an organismal wide label for proliferative zones, and an endogenous cell cycle reporter in later sections.

#### Intra-cellular trafficking

To generate a reporter line allowing monitoring sub-cellular trafficking of endosomes and exosomes, we targeted Rab11a (Ras-Related Protein), a small GTPase and known marker of intra-cellular trafficking organelles in vertebrates (Welz et al., 2014, Cullen and Steinberg, 2018, Stenmark, 2009). We generated an N-terminus tagged *eGFP-rab11a* allele that labels sub-cellular trafficking organelles (Figure 2C, Figure S5 and Supplementary Movies 2/3/4). As a proof of principle, we detected high levels of *eGFP-rab11a* in cells of the spinal cord (Figure 2C yellow arrowhead) and in neuromasts of the lateral line (Figure 2C magenta arrowhead, Figure S5, Supplementary Movie 4, n=6 embryos). Using the *eGFP-rab11a* KI line, we were also able to observe dynamics of intra-cellular organelle trafficking *in vivo* both in individual skin epithelial cells in the mid-trunk region and in the caudal fin region of developing medaka embryos (Supplementary Movies 2/3, n=4) validating the utility of this line as a sub-cellular trafficking marker in medaka.

#### Stress granule marker

We were able to generate a *g3b1-eGFP* KI line by targeting *eGFP* to the 11th exon of the medaka *g3bp1* gene. G3bp1 (GTPase activating protein SH3-domain binding protein) is a DNA/RNA-binding protein and an initiating factor involved in stress granule formation (Irvine et al., 2004, Yang et al., 2020). Stress granules are non-membrane bound cell compartments, which form under cellular stress and accumulate non-translating mRNA and protein complexes, and play an important role in cellular protection by regulating mRNA translation and stability (Decker and Parker, 2012, Protter and Parker, 2016). Under normal conditions G3bp1-eGFP is expressed in the cytoplasm (Figure 2F, Supplementary Movie 5, n=8 embryos) but upon stress (temperature shock), we observe that the protein changes its localization and accumulates in cytoplasmic foci corresponding to forming stress granules (Figure 2F’ yellow arrowheads, Supplementary Movie 5, n=8 embryos). This is consistent with previous reports showing similar changes in the localization of G3bp1 in response to stress in a number of organisms (Guarino et al., 2019, Wheeler et al., 2016, Kuo et al., 2020). The initial characterization of the *g3bp1-eGFP* line shows its potential to serve as a real-time *in vivo* reporter for the dynamics of stress granules formation in a vertebrate model.

#### Muscle cell marker

To label muscle cells, we targeted muscular *myosin heavy chain* with *mNeonGreen.* Myosins are a highly conserved class of motor proteins implicated in actin microfilament reorganization and movement (Sellers, 2000, Hartman and Spudich, 2012). We generated an N-terminus fusion of *mNeonGreen-myosinhc* KI that exclusively labels muscle cells (Figures 1D/2D and Figure S4, n>10 embryos). In the medaka myotome, we were able to observe *mNeonGreen-myosinhc* chains of individual sarcomeres (A-bands separated by the I-bands), indicating that tagged Myosinhc is incorporated correctly in muscle fibers (Taylor et al., 2015, Loison et al., 2018). We use this line to record the endogenous dynamics of Myosinhc during muscle growth *in vivo* for the first time to the best of our knowledge, in a vertebrate model (Supplementary Movie 6 n=9 embryos). The *mNeonGreen-myosinhc* line therefore enables the *in vivo* recording of endogenous Myosinhc dynamics during myogenesis in medaka.

#### Cell adhesion marker

Cadherins are a highly conserved class of trans-membrane proteins that are essential components of cell-cell adhesion and are thus expressed on cellular membranes (Leckband and de Rooij, 2014). A large number of *cadherin* genes exist in vertebrates where they exhibit tissue specific expression patterns and are implicated in various developmental processes (Halbleib and Nelson, 2006). We decided to tag the C-terminus of medaka *cadherin 2 (cdh2, n-cadherin*) with *eGFP*. *cdh2 is* known to be expressed primarily in neuronal tissues in a number of vertebrates (Harrington et al., 2007, Suzuki and Takeichi, 2008). The *cdh2-eGFP* KI line shows cellular membrane expression in a variety of neuronal and non-neuronal tissues including the spinal cord, the notochord (Figure 2E, n=5 embryos) and neuromasts of the lateral line (Figure S5, Supplementary Movie 7, n=5 embryos), in addition to the developing heart (data not shown) (Chopra et al., 2011). The high expression of *cdh2* in both differentiated notochord cell types (Figure 2E, Figure S5) has not been previously reported in medaka but is not unexpected as this tissue experiences a high level of mechanical stress and requires strong cell-cell adhesion (Lim et al., 2017, Adams et al., 1990, Garcia et al., 2017, Seleit et al., 2020). The *cdh2-eGFP* KI is thus the first endogenously tagged cadherin family member in teleosts and can be used to study dynamics of *n-cadherin* distribution *in vivo* during vertebrate embryogenesis (Supplementary Movie 8).

### *mScarlet-pcna:* an organismal-wide marker for proliferative zones

We reasoned that the novel *mScarlet-pcna* line can act as an organismal-wide *bona fide* marker for the location of proliferative cells within any tissue or organ of interest. We therefore decided to generate double transgenic animals with *eGFP-cbx1b* as a ubiquitous nuclear marker and *mScarlet-pcna* as a label for cycling cells (Figure 3A-C’’). As a proof of principle, we set out to investigate the location of proliferative zones in a number of organs and tissues in medaka. We began by assessing the position of proliferative cells in neuromast organs of the lateral line (Seleit et al., 2017b, Pinto-Teixeira et al., 2015, Romero-Carvajal et al., 2015). Neuromasts are small rosette shaped sensory organs located on the surface of teleost fish that sense the direction of water flow and relay the information back to the Central Nervous System (CNS) (Seleit et al., 2017a, Romero-Carvajal et al., 2015, Jones and Corwin, 1993, Wada et al., 2013). They consist of four cell types: differentiated Hair Cells (HCs) in the very center, underlying Support Cells (SCs), a ring of Mantle Cells (MCs) and neuromast Border Cells (nBCs) (Seleit et al., 2017b, Dufourcq et al., 2006). Previous work in medaka has established MCs to be the true life-long neural stem cells within mature neuromast organs (Seleit et al., 2017b). While the *eGFP-cbx1b* labels all neural cells within a mature neuromast organ (HCs, SCs and MCs) (Figure 3A), *mScarlet-pcna* expression matches the previously reported location of proliferative MCs (Seleit et al., 2017b) (Figure 3A’-A’’, white arrowhead). Neither the differentiated HCs nor the SCs directly surrounding them show evidence of Pcna expression in mature neuromast organs under homeostatic conditions in medaka (Figure 3A-A’’, n=10 neuromast organs). Our results validate the utility of *mScarlet-pcna* as an *in vivo* marker of proliferative cells. Previous work has shown that nBCs are induced to form from epithelial cells that come into contact with neuromast precursors during organ formation and that these induced cells become the stem cell niche of mature neuromast organs (Seleit et al., 2017b). However, an open question is whether transformed nBCs are differentiated, post-mitotic cells or whether they remain cycling. Utilizing the *mScarlet-pcna* line we were able to observe nBCs (4/42) in late S-phase of the cell cycle, as evident by the presence of nuclear speckles, in mature neuromast organs (Figure 3A’-A’’, yellow arrowheads). This provides direct evidence that nBCs retain the ability to divide and are thus not post-mitotic cells.

**Figure 3:**
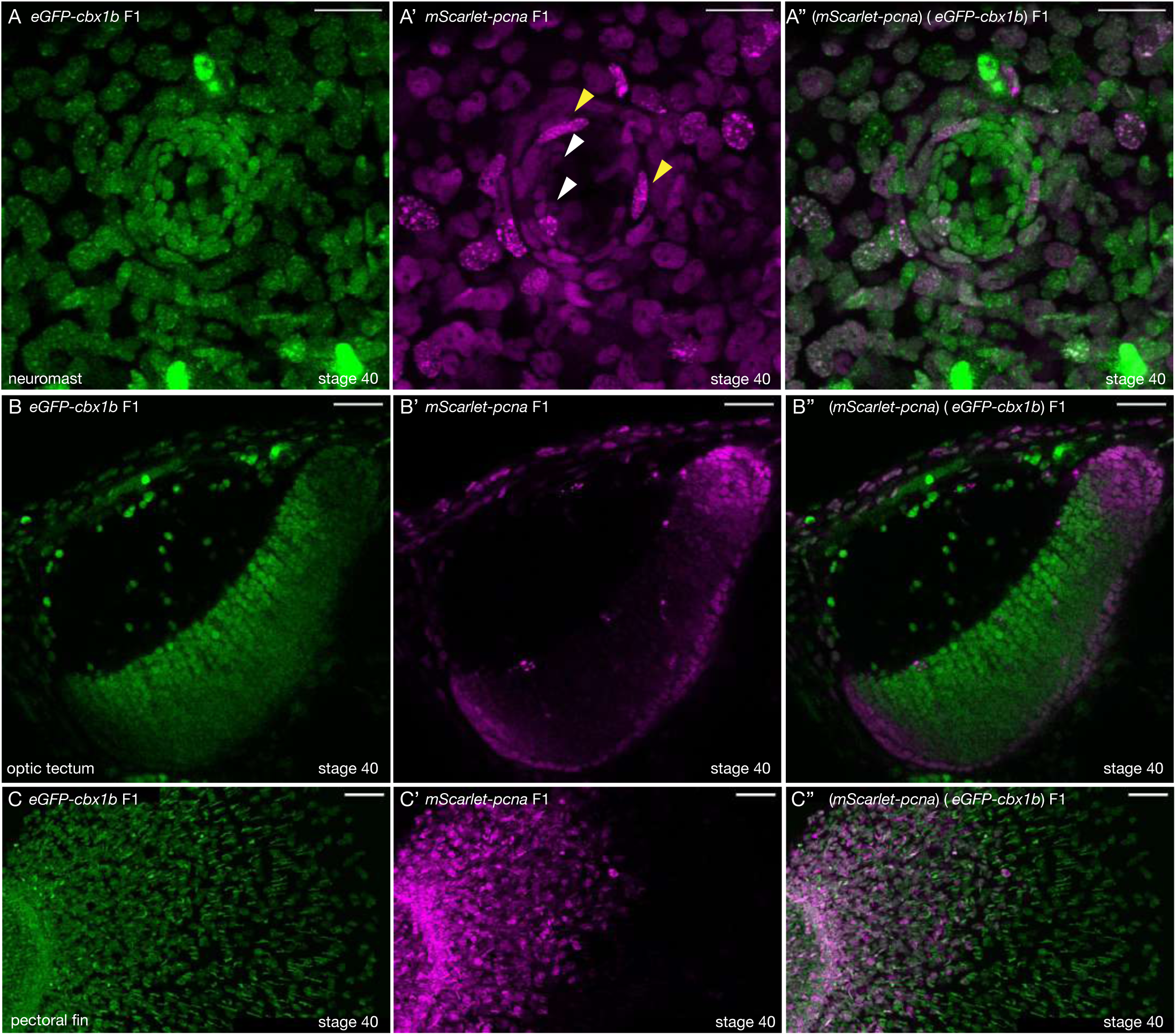
*mScarlet-pcna* line acts as an organismal-wide marker for proliferative zones. (**A-A’’**) (*eGFP-cbx1b*) (*mScarlet-pcna*) double positive stage 40 medaka embryo. Maximum projection of a mature secondary neuromast (center of image) within the lateral line system. surrounded by epithelial cells labelled by: endogenous eGFP-Cbx1b in (A) and endogenous mScarlet-Pcna in (A’). The merge is shown in (A’’). (A) eGFP-Cbx1b is a ubiquitous nuclear marker and labels all cell types within a mature neuromast. Those are: Hair Cells (HCs) in the center of a neuromast, surrounded by Support Cells (SCs) and an outer ring of Mantle Cells (MCs) surrounded by the elongated neuromast Border cells (nBCs). (A’) mScarlet-Pcna labels cycling cells, which are located at the very edge of the mature neuromast organ, a proportion of Mantle Cells (MCs) express mScarlet-Pcna (white arrowheads). Neuromast border cells (nBCs) also express mScarlet-Pcna. Speckles can be seen in several mScarlet-Pcna *positive* nBC nuclei (yellow arrowheads), indicating cells in late S phase of the cell cycle. (A’’) Merged image. n= 10 neuromast organs. Scale bar = 20µm. (**B-B’’**) (*eGFP-cbx1b*) (*mScarlet-pcna*) stage 40 medaka embryo. Single Z-slice showing the medaka optic tectum. (B) eGFP-Cbx1b is a ubiquitous nuclear marker (B’) whereas mScarlet-Pcna labels a subset of cells at the outer periphery of the optic tectum, indicating the position of proliferative cells in this tissue. A graded expression of mScarlet-Pcna is observed, with more central cells in the optic tectum losing the expression of mScarlet-Pcna. (B’’) Merged image. n= 4 embryos. Scale bar = 30µm. (**C-C’’**) Maximum projection of the pectoral fin of stage 40 medaka embryos. (C) eGFP-Cbx1b is a ubiquitous nuclear marker (C’) while a subset of cells is labeled by mScarlet-Pcna indicating the position of proliferative cells. Note the proximal to distal gradient of mScarlet-Pcna expression, with proliferative cells at the base of the fin (left) and differentiated cells at the outer edges of the fin (right) (C’’) Merged image. n=4 embryos. Scale bar = 50µm.

Next, we turned our attention to the optic tectum, which is essential for integrating visuomotor cues in all vertebrates (Lavker and Sun, 2003, Alunni et al., 2010, Nguyen et al., 1999). We show that proliferative cells in the optic tectum of medaka are located at the lateral, caudal and medial edge of the tectum in a crescent-like topology (Figure 3B-B’’). Moreover, *mScarlet-pcna* expression is graded, with the more central cells gradually losing expression of Pcna (Figure 3B’, n=4 embryos). This is in line with previous histological findings using BrdU stainings in similarly staged medaka embryos (Nguyen et al., 1999, Alunni et al., 2010). We next analyzed the expression of *mScarlet-pcna* in the developing pectoral fin (Figure 3C-C’’ n=4 embryos). We found that cells located proximally expressed the highest levels of *mScarlet-pcna,* with *mScarlet-pcna* expression decreasing gradually along the proximo-distal axis. To the best of our knowledge this proliferation pattern has not been previously reported and our data provides evidence that the differentiation axis of the pectoral fin is spatially organized from proximal to distal in medaka. Lastly, we reveal that proliferative cells are present in the spinal cord of stage 40 medaka embryos, a finding that has not been previously reported, and we show that these *mscarlet-pcna* positive cells occur in clusters preferentially located on the dorsal side of the spine (Figure S6, n=4). The newly developed *mScarlet-pcna* therefore acts as a stable label of proliferative cells and as such can be used to uncover the location of proliferation zones *in vivo* within organs or tissues of interest in medaka.

### *mScarlet-pcna:* an endogenous cell cycle reporter

In addition to its use as a marker for cells in S-phase, it has been shown that endogenously-tagged Pcna can be used to determine all other cell cycle phases. This is based on the fact that both the levels and dynamic distribution of Pcna shows reproducible characteristics in each phase of the cell cycle (Held et al., 2010, Piwko et al., 2010, Santos et al., 2015, Zerjatke et al., 2017). To assess whether the endogenous *mScarlet-Pcna* line recapitulates these known characteristic expression features during the cell cycle, we aimed to quantitatively analyze endogenous *mScarlet-Pcna* levels in individual cells during their cell cycle progression. To this end, we imaged skin epithelial cells located in the mid-trunk region of medaka embryos (Figure 4A-D, Figures S7/S8). Cells in the G1 phase of the cell cycle have been shown to decrease the levels of Pcna within the nucleus over time (Figure S7, Supplementary Movie 9, n= 9 epithelial cells) (Zerjatke et al., 2017). On the other hand, cells progressing through to S phase have been shown to increase the levels of Pcna expression within the nucleus over time (Leonhardt et al., 2000, Piwko et al., 2010, Leung et al., 2011, Santos et al., 2015, Barr et al., 2016, Zerjatke et al., 2017, Held et al., 2010). All the tracked epithelial cells that eventually underwent a cellular division during our time-lapse imaging showed an increase in nuclear intensity of mScarlet-Pcna over time (Figure 4A-B, n= 9 epithelial cells). Late S phase is categorized by the occurrence of nuclear speckles of Pcna marking the presence of replication foci (Leonhardt et al., 2000, Piwko et al., 2010, Leung et al., 2011, Santos et al., 2015, Barr et al., 2016, Zerjatke et al., 2017), which we observe in every dividing epithelial cell prior to cell division (Figure 4A, Supplementary Movie 10, n= 9 epithelial cells). The S/G2 transition is marked as the point of peak pixel intensity distribution of endogenous Pcna within the nucleus (Zerjatke et al., 2017), which we are able to obtain for each dividing cell by a combination of 3D surface plots and histograms of pixel intensity distributions over time (Figure 4C-C’’’, Figures S7/S8 and Supplementary Movies 10/11/12, n= 9 epithelial cells). While the M phase is marked with a sharp decrease in nuclear levels of Pcna (Zerjatke et al., 2017), which can be consistently seen in the endogenous *mScarlet-pcna* intensity tracks of epithelial cells undergoing division (Figure 4B, Figures S7/S8, Supplementary Movies 10/11/12, n= 9 epithelial cells). Cells are in the G2 phase of the cell cycle in the time frame between the S/G2 transition point to the beginning of M phase. We therefore provide first evidence that the *mScarlet-pcna* line recapitulates known dynamics of Pcna within the nucleus (Held et al., 2010, Piwko et al., 2010, Santos et al., 2015, Zerjatke et al., 2017) and that it can therefore be utilized as an endogenous ‘all-in one’ cell cycle reporter in vertebrates.

**Figure 4:**
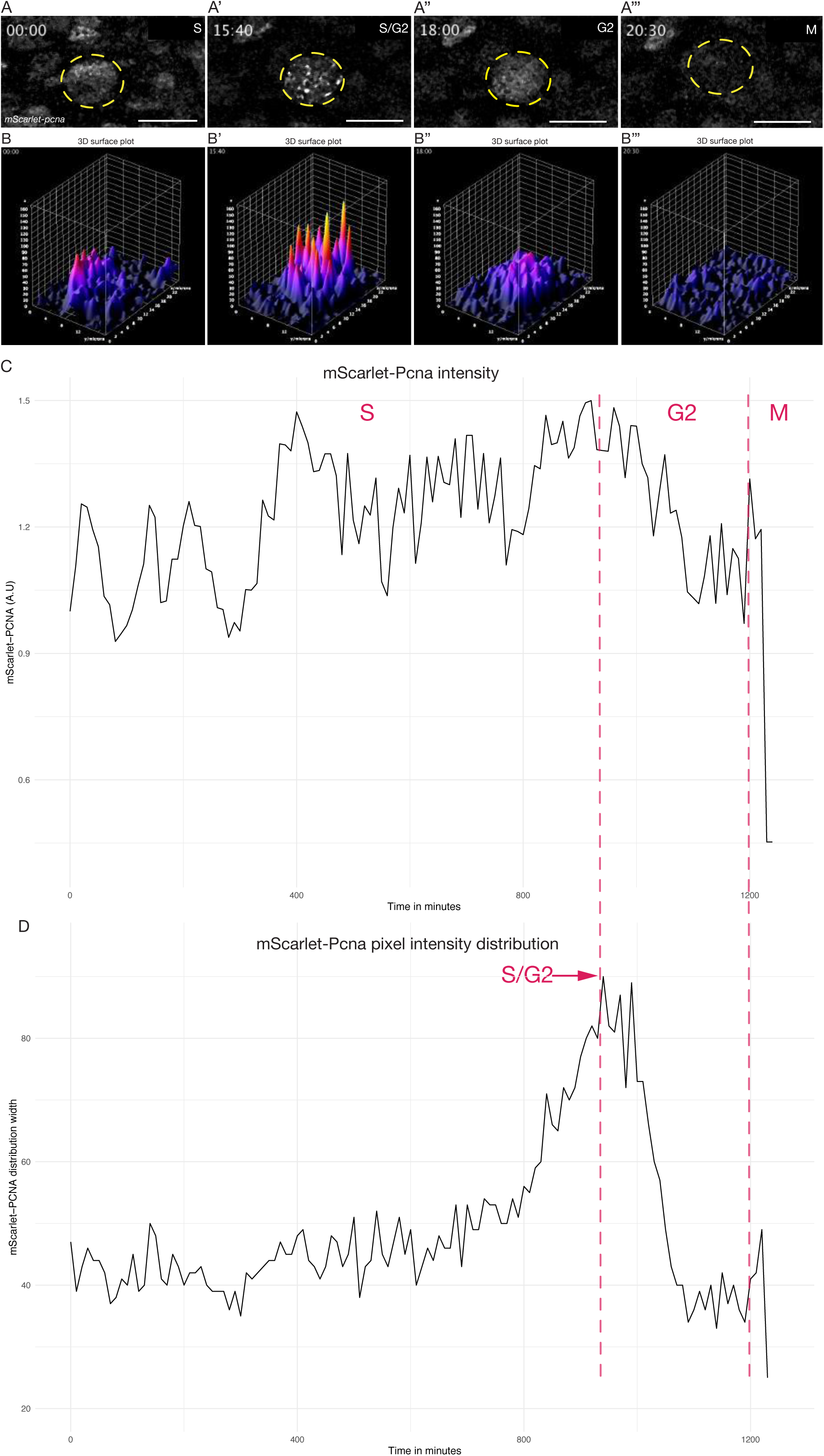
Quantitative live cell-tracking of endogenous *mScarlet-Pcna* levels enables cell cycle phase classification. (**A-A’’’**) Selected frames from a time-lapse imaging of a mScarlet-Pcna positive skin epidermal cell nucleus (yellow circle) undergoing cell division. The different phases of the cell cycle are deduced from mScarlet-Pcna expression as highlighted within the panels. Late S phase can be distinguished by the presence of nuclear speckles that correspond to replication foci. n= 9 skin epithelial cells. Time in hours. (**B-B’’’**) 3D surface plots of cell from (A). The S/G2 transition is marked as the point of peak pixel intensity distribution within the nucleus, which is reached at 15:40h (B’) and is equivalent to the largest width of mscarlet-Pcna pixel intensity distribution shown in panel D (red arrow). n= 9 skin epidermal cells. (**C**) Normalized mScarlet-Pcna intensity within the nucleus from cell in (A) over the course of one cell division. Vertical dashed red lines demarcate the different cell cycle phases based on the intensity and dynamic distribution of mScarlet-Pcna within the nucleus. Initially an increase of endogenous mScarlet-Pcna expression over time indicates cells in S phase of the cell cycle. M phase is characterized by a sharp drop in nuclear mScarlet-Pcna levels begining at 20:00h. (**D**) Width of pixel intensity distribution over time on cell from (A). The S/G2 transition is marked as the point of peak pixel intensity distribution within the nucleus, which is reached at 15:40h and is equivalent to the largest width of mscarlet-Pcna pixel intensity distribution (red arrow). n= 9 skin epidermal cells.

## Discussion

Despite the CRISPR/Cas9 system being repurposed as a broad utility genome editing tool almost a decade ago (Jinek et al., 2012, Cong et al., 2013) and despite its revolutionary impact as a method to generate knock-ins by homology directed repair (Wang et al., 2016, Danner et al., 2017, Jasin and Haber, 2016, Ceccaldi et al., 2016), there is still a paucity of precise, single-copy fusion protein lines in vertebrates, in general, and in teleost fish in particular. In fact, in medaka there are a total of three validated single-copy fusion protein lines reported prior to this work (Gutierrez-Triana et al., 2018) while in zebrafish only a handful of lines have been generated (Wierson et al., 2020, Hisano et al., 2015, Hoshijima et al., 2016, Shin et al., 2014, Wierson et al., 2019). This underscores the complexity of generating and validating precise single-copy fusion protein knock-in lines in teleost models. Previous techniques to generate large KIs (such as fluorescent reporters) required the usage of plasmid vectors commonly containing long homology arms (>200bp) (Zu et al., 2013, Auer and Del Bene, 2014, Shin et al., 2014). Problems arising during and after injection include DNA concatemerization of the donor construct (Gutierrez-Triana et al., 2018, Auer et al., 2014, Winkler et al., 1991), in addition to possible imprecise and off-target integration of either the fluorescent protein sequence or the plasmid backbone (Auer et al., 2014, Gutierrez-Triana et al., 2018, Won and Dawid, 2017, Wierson et al., 2020, Cristea et al., 2013, Shin et al., 2014, Wierson et al., 2019, Yao et al., 2017). The vast majority of reported HDR mediated knock-ins in teleosts rely on *in vivo* linearization of the plasmid donors. This strategy is utilized due to the observation that, although linear dsDNA donors can drive HDR, they might be prone to degradation, concatemerization, and are generally thought to be more toxic than plasmid donors (Auer et al., 2014, Cristea et al., 2013, Shin et al., 2014, Ota et al., 2014). Plasmid donors therefore contain an additional guide RNA sequence to drive *in vivo* linearization in order to synchronize the availability of the linear DNA donor with Cas9 activity (Auer et al., 2014, Cristea et al., 2013, Shin et al., 2014, Lisby and Rothstein, 2004, Ota et al., 2014). We reasoned that directly injecting PCR amplified linear DNA with short homology arms (∼35bp) could be highly effective since these donors are relatively small (∼780 bp) compared to plasmids (several kbs), and therefore a small quantity of donors (∼10 ng/ul) will provide a large number of molecules (∼20 nM) available to engage the HDR machinery following the Cas9 induced DSB. Building on recent improvements in CRISPR/Cas9 knock-in strategies, we used 5’ biotinylated primers in order to limit *in vivo* concatemerization of the donor construct (Gutierrez-Triana et al., 2018), and synthetic sgRNAs were used to increase the efficiency of DSBs by Cas9 (Paix et al., 2015, Kroll et al., 2021, Hoshijima et al., 2019, Li et al., 2019). In addition, we utilized a monomeric streptavidin tagged Cas9 that has a high affinity to the biotinylated donor fragments to increase targeting efficiency (Gu et al., 2018). We report that this approach is a highly efficient, precise and scalable strategy for generating single-copy fusion proteins (Table1 and Table S1). Since the repair donors are synthesized by PCR amplification, we eliminate the need for both cloning and a second gRNA for *in vivo* linearization. Very recently a similar approach to the one we present here showed the potential to generate KI lines in zebrafish by targeting non-coding regions with PCR amplified donor constructs (Levic et al., 2021). All in all, the strategy we utilize significantly simplifies the process of endogenous protein tagging in a vertebrate model.

An important aspect of any knock-in strategy to generate fusion proteins is its precision. The validation process of single copy insertions is complicated in approaches that use long homology arms (>200bps) to generate knock-ins as concatemerization cannot be easily ruled out. *Locus* genotyping by PCR and Sanger sequencing is difficult when using primers external to the repair donor due to the large size of the expected fragment. Internal primers within the donor (junction-PCR) have been used to avoid this limitation, but this can lead to PCR artefacts and crucially, it does not rule out concatemerization of the injected dsDNA (Won and Dawid, 2017, Gutierrez-Triana et al., 2018, Wierson et al., 2020). Southern blotting is considered the gold standard to assess single-copy integration (Wierson et al., 2020, Gutierrez-Triana et al., 2018, Won and Dawid, 2017). While it has its advantages, Southern Blotting depends on experimental design (genomic DNA digestion strategy) and probe design/sensitivity, and therefore cannot exclude that part of the donor construct or part of the vector backbone integrates elsewhere in the genome. Indeed, it has been reported that plasmid donors can lead to additional unwanted insertions in the genome (Won and Dawid, 2017, Wierson et al., 2020). We address those issues by performing WGS with high coverage on knock-in lines and provide evidence that our approach yields single-copy integration only at the desired *locus* (Figure S2). In addition, utilizing repair donors with short homology arms on both ends (30-40bp) simplifies the validation of the insertion by using primers that sit outside the targeting donor fragment. These external primers can then be used for genotyping of the full insertion by simple PCR followed by Sanger sequencing to know the precise nature of the edit (Figure S2). We show that the usage of donor fragments with short homology arms, in combination with high coverage WGS, to be important aspects in validating the precision of single-copy CRISPR/Cas9 mediated knock-in lines in vertebrate models.

We were able to generate six novel endogenous protein fusion lines that significantly expand the repertoire of genetic tools to track cellular dynamics in medaka. The *eGFP-cbx1b* KI line is the first reported endogenous ubiquitous nuclear marker in teleosts (Nielsen et al., 2001, Lomberk et al., 2006, Gilmore et al., 2016). The generation of truly ubiquitous lines by transgene over-expression in teleost fish (Centanin et al., 2014, Burket et al., 2008) is a difficult endeavor and requires constant monitoring for variegation and silencing (Goll et al., 2009, Akitake et al., 2011, Burket et al., 2008, Stuart et al., 1990). Yet these ubiquitous fluorescent reporter lines are invaluable tools for researchers. Ubiquitous fusion-proteins expressed from the endogenous *locus* avoid potential issues with transgene over-expression and variegation. The highly conserved *cbx1b locus* could therefore provide an alternative strategy to generate faithful ubiquitous nuclear markers in other teleosts and non-model organisms. In addition, this *locus* could serve as a landing site for ubiquitous expression of genetic constructs (for *e.g.* utilizing a T2A peptide) in medaka (Li et al., 2019, Kim et al., 2011). Next, we validate the use of g*3bp1-eGFP* knock-in as a stress granule formation marker, and utilizing 4-D live-imaging show the formation of stress granules in response to temperature shock in real-time, as previously shown in other models using a variety of stress conditions(Guarino et al., 2019, Kuo et al., 2020, Wheeler et al., 2016, Decker and Parker, 2012, Protter and Parker, 2016). This line can therefore be used both as an *in vivo* marker of stress conditions and to study the process of stress granule formation. The *eGFP-rab11a line* serves as an intra-cellular trafficking (Welz et al., 2014, Cullen and Steinberg, 2018) marker that allows us to dynamically follow exosomes and endosomes *in vivo*. We report that both neuromasts and the spinal cord show substantially higher expression of *rab11a* than other tissues, the basis of this remains unclear but could indicate that these tissues exhibit higher levels of protein turn-over. Despite being a highly conserved protein involved in myogenesis (Sellers, 2000, Hartman and Spudich, 2012), no endogenous KI of any myosin family member has been reported in teleosts. The *mNeonGreen-myosinhc* knock-in enables the detection and recording of endogenous myosin dynamics *in vivo* during muscle growth in a vertebrate model. We also generate *cdh2-eGFP* as the first reported fusion-protein line for a Cadherin family member in teleosts (Leckband and de Rooij, 2014, Halbleib and Nelson, 2006) and show that it is primarily expressed in neuronal tissues including the spinal cord and neuromasts. Since N-cadherin has been shown to be involved in Epithelial–Mesenchymal Transition (EMT) (Harrington et al., 2007, Suzuki and Takeichi, 2008, Desclozeaux et al., 2008), this line can be used to study dynamical changes in N-cadherin distribution *in vivo* facilitating our understanding of EMT and other fundamental cell adhesion processes in vertebrates. Lastly, we generate and characterize the *mScarlet-pcna* knock-in line and discuss its usage and implications across teleosts below.

An overarching goal of developmental and stem cell biology is to discover the location of stem and progenitor cells in different organs and tissues, followed by a molecular characterization of their properties (Rhee et al., 2006, Nowak et al., 2008, Snippert et al., 2010, Buczacki et al., 2013, Lu et al., 2012, Lavker and Sun, 2003). Major advances have relied on finding resident stem cell markers that differentiates stem cells from other cell types within the same tissue, followed by BrdU/IdU staining to confirm their proliferative abilities (Nguyen et al., 1999, Rhee et al., 2006, Nowak et al., 2008, Nowak and Fuchs, 2009, Alunni et al., 2010, Snippert et al., 2010, Lu et al., 2012, Buczacki et al., 2013, Stolper et al., 2019, Tsingos et al., 2019). However, BrdU/IdU staining requires the sacrifice of the animal precluding the ability to perform 4D live-imaging to analyze stem cell behavior *in vivo* over time. The medaka knock-in line with endogenously labeled Pcna that we present here helps to circumvent this limitation. In addition, since Pcna is expressed exclusively in cycling cells (Yamaguchi et al., 1995, Thacker et al., 2003, Buttitta et al., 2010, Zerjatke et al., 2017), it has the potential to be used to discover the location of proliferative zones *in vivo* within any organ or tissue of interest. We provide proof of principle evidence that the *mScarlet-pcna* KI line acts as a *bona fide* marker for proliferative zones in a variety of tissues in medaka fish. This line therefore represents an important new tool for stem cell research in medaka. A similar strategy could be adopted to generate endogenously tagged Pcna both in the teleost field and in other organisms.

In addition to its use as a *bona fide* marker for proliferative zones, we provide evidence that the *mScarlet-pcna* line can be used as an endogenous cell cycle reporter in medaka. It has previously been shown that both the levels and dynamic distribution of Pcna are indicative of the different the cell cycle phases (Held et al., 2010, Piwko et al., 2010, Santos et al., 2015, Zerjatke et al., 2017). This led researchers to successfully utilize it as an ‘all-in-one’ cell cycle reporter in mammalian cells (Held et al., 2010, Piwko et al., 2010, Santos et al., 2015, Zerjatke et al., 2017). By quantitatively tracking endogenous Pcna levels during one cell cycle in epidermal cells of medaka fish, we were able to confirm the dynamic nature of *mScarlet-pcna* expression, which correlated with the previously described behavior of the Pcna protein within the nucleus of other vertebrates (Held et al., 2010, Piwko et al., 2010, Santos et al., 2015, Zerjatke et al., 2017). As such, we provide proof of principle evidence that the *mScarlet-pcna* line can be successfully used as an endogenous cell cycle reporter in a teleost model. Using the visualization of endogenous Pcna for cell cycle phase classification offers an attractive alternative to cell cycle reporters that rely on the insertion of two-colour transgenes, such as the FUCCI system (Sugiyama et al., 2009, Dolfi et al., 2019, Araujo et al., 2016, Bajar et al., 2016, Oki et al., 2014, Sakaue-Sawano et al., 2008). First, by using endogenous fusion proteins there is no requirement for over-expression of cell cycle regulators. Second, the potential issue with transgene variegation and silencing is avoided (Akitake et al., 2011, Goll et al., 2009, Burket et al., 2008, Stuart et al., 1990). Finally, utilizing a single color cell-cycle reporter allows its simultaneous use with other fluorescent reporters during live-imaging experiments. Due to the high conservation of Pcna in eukaryotes, developing Pcna reporters in other model organisms using a similar strategy is an attractive possibility to pursue.

---

**Supplementary Figure 1:**
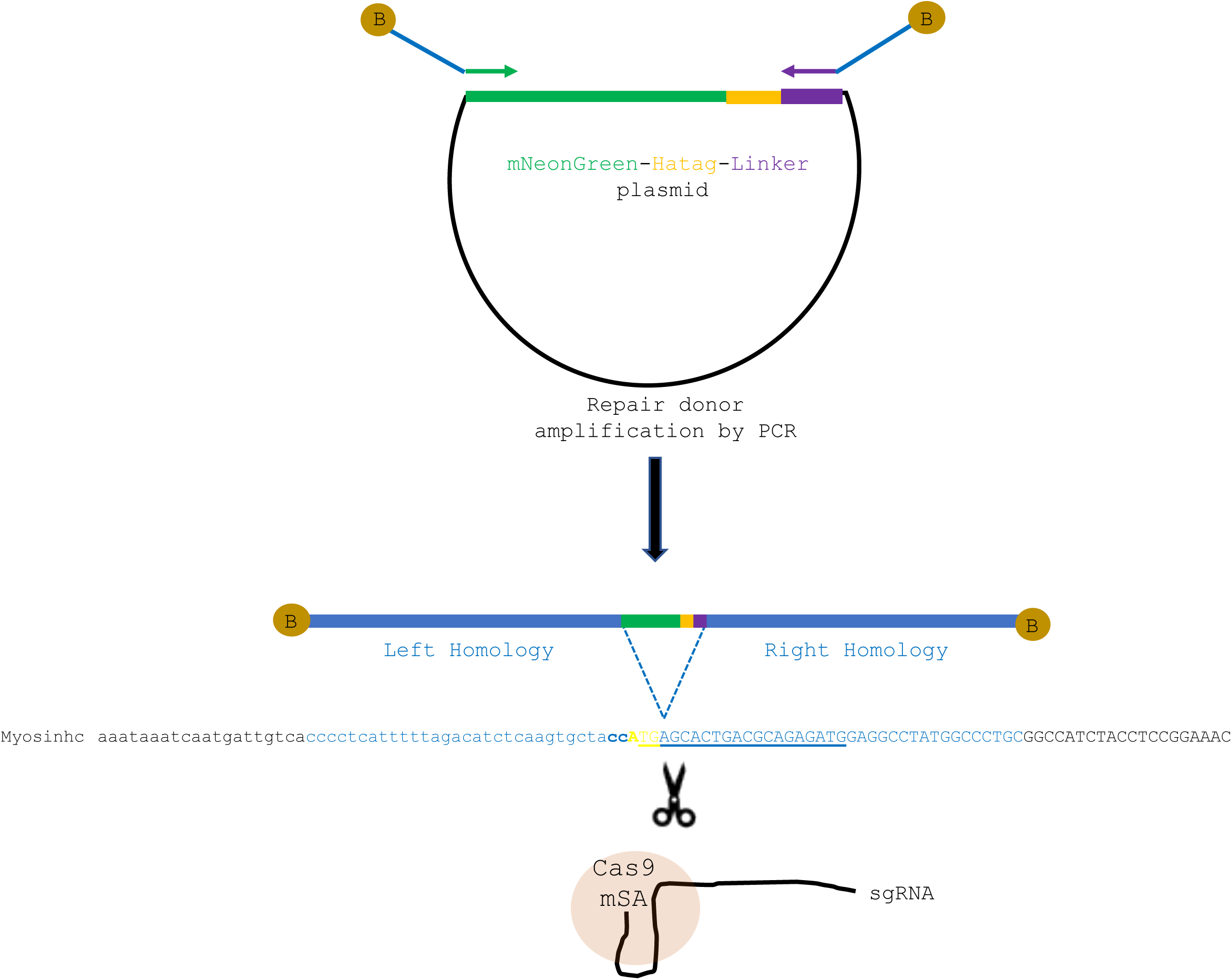
Schematic representation of *mNeonGreen-myosinhc* tagging strategy. (**A**) PCR amplification of *mNeonGreen-HAtag-Linker* (Green, Orange and Purple) using primers homologous to the extremity of the insert and containing flanking sequence corresponding to the homology arms (Blue) for insertion at the ATG of *myosinhc* gene (Yellow). The primers contain 5’end Biotins (Brown circle). (**B**) Insertion of *mNeonGreen-HAtag-Linker* just downstream the ATG of *myosinhc* (Yellow) following Cas9-mSA / sgRNA DNA induced cut. Lower letters denote noncoding sequence, upper letters coding sequence, bold the sgRNA PAM, underlined the sequence upstream the PAM recognized by the sgRNA, blue letters the sequence homologous between the *myosinhc locus* and the PCR donor.

**Supplementary Figure 2.**
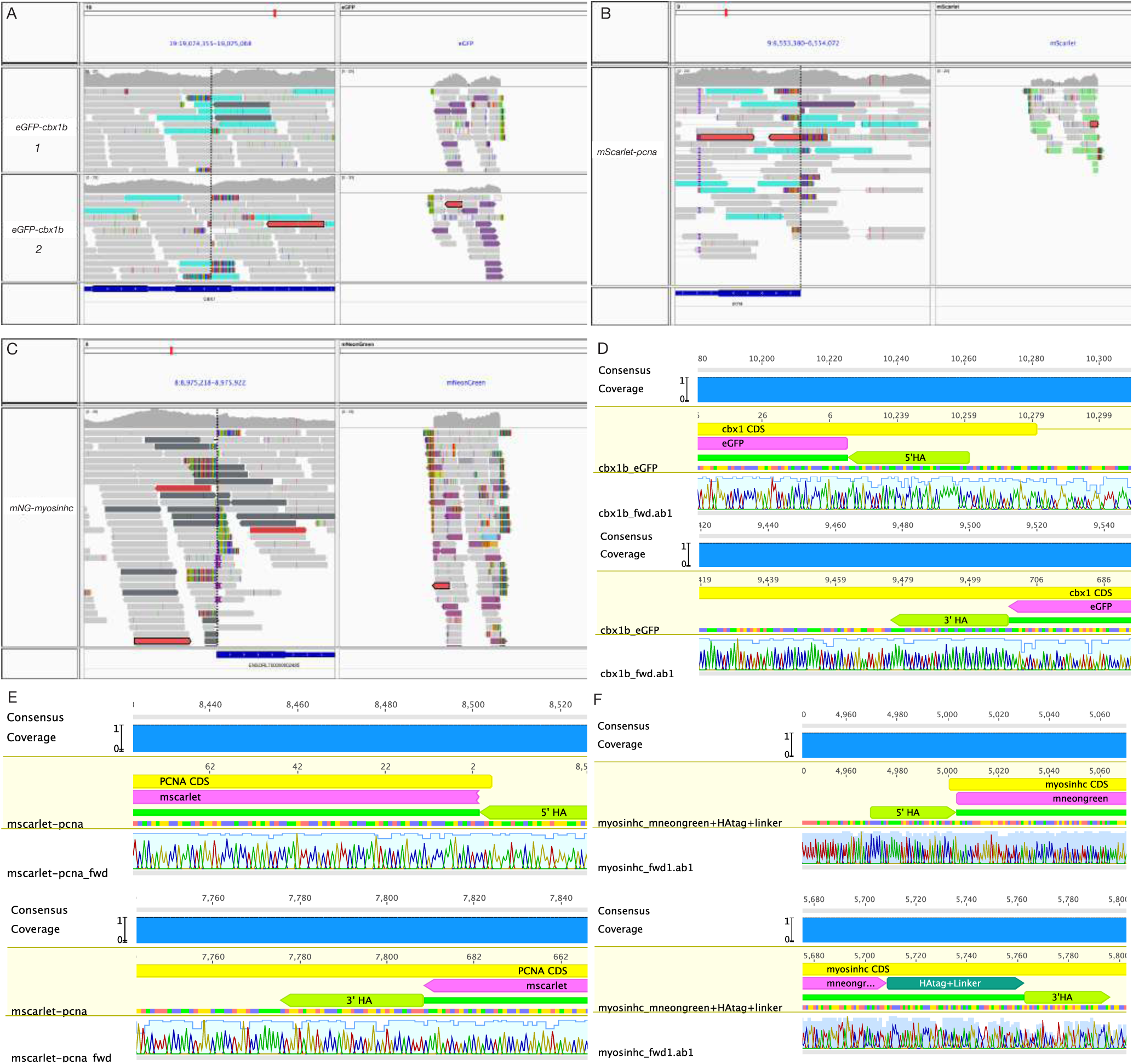
Alignment of whole genome sequencing reads (WGS) to fluorescent protein sequences: (**A**) *eGFP* integration in *cbx1b locus*. Paired-end sequenced reads from *eGFP-cbx1b* F1 embryos, 2 biological replicates (gDNA1 and gDNA2) from the same F0 founder are shown in the upper and lower panel, coloured in grey for concordant mappings and coloured in turquoise/purple for inter-chromosomal paired-ends with one mate mapped to chr19 (left panel) and one mate mapped to *eGFP* (right panel). The predicted integration site is shown as a vertical dashed line with soft-clipped reads (shown as coloured bases) to the left and right of the integration site. Above the sequenced reads is the basepair-level coverage histogram. Coverage of *eGFP-cbx1b_gDNA1* is 20.4X and for *eGFP-cbx1b_gDNA2* is 23.6X. (**B**) *mScarlet* integration in *pcna locus*. Paired-end sequenced reads from *mScarlet-pcna* F1 embryos, coloured in grey for concordant mappings and coloured in turquoise/green for inter-chromosomal paired-ends with one mate mapped to chr9 (left panel) and one mate mapped to *mScarlet* (right panel). The predicted integration site is shown as a vertical dashed line with soft-clipped reads (shown as coloured bases) to the left and right of the integration site. Above the sequenced reads is the basepair-level coverage histogram. Coverage of *mScarlet-pcna* is 14.4X. (**C**) *mNeongreen* integration in *myosinhc locus*: Paired-end sequenced reads from *mNG-myosinhc* F1 embryos, coloured in grey for concordant mappings and coloured in dark grey/purple for inter-chromosomal paired-ends with one mate mapped to chr8 (left panel) and one mate mapped to *mNeonGreen* (right panel). The predicted integration site is shown as a vertical dashed line with soft-clipped reads (shown as coloured bases) to the left and right of the integration site. Above the sequenced reads is the basepair-level coverage histogram. Paired-end analysis yielded a second, weakly supported insertion of *mNeonGreen* at chr12:9083923 which could not be confirmed by PCR, suggesting that this is a false positive prediction or a mosaic insertion of very low frequency. Coverage of *mNG-myosinhc* is 14.5X. (**D**) Sanger sequencing read from *eGFP-cbx1b* F1 fin-clipped adult shows scarless integration of *eGFP* into the *cbx1b locus* in both 5’ and 3’ junctions (**E**) Sanger sequencing reads from *mScarlet-pcna* F1 fin-clipped adult shows scarless integration of *mScarlet* into the *pcna locus* at the 3’ junction. While the 5’ junction shows correct in-frame fusion of endogenous *pcna* to mscarlet, we were able to detect a partial duplication of the 5’ homology arm that occured 22bp upstream of the start codon of endogenous *pcna* (for details see materials and methods). (**F**) Sanger sequencing reads from *mNG-myosinhc* F1 fin-clipped adult shows scarless integration of *mNeonGreen* into the *myosinhc locus* in both 5’ and 3’ junctions.

**Supplementary Figure 3:**
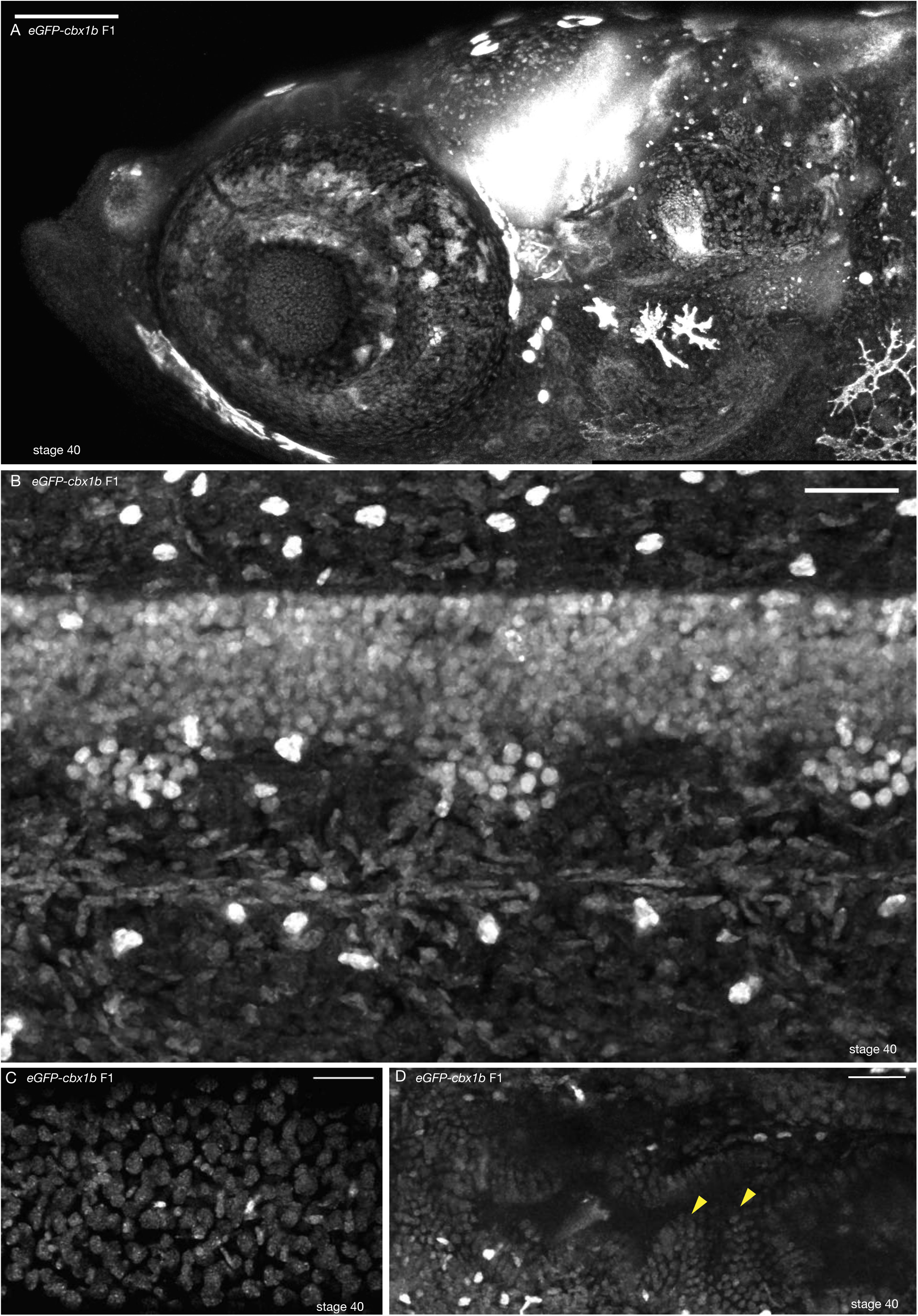
Ubiquitous nuclear expression of *eGFP-cbx1b*. (**A**) Anterior head region of *eGFP-cbx1b* F1 stage 40 medaka embryo. eGFP-Cbx1b is expressed ubiquitously with a nuclear localization. n>10 embryos. Scale bar = 100µm. (**B**) Maximum projection of spinal cord and notochord in the mid-trunk region of *eGFP-cbx1b* stage 40 medaka embryo. All nuclei are labeled by eGFP-Cbx1b. n>10 embryos. Scale bar = 30µm (**C**) A subset of epithelial cell nuclei positive labeled by eGFP-Cbx1b. n>10 embryos. Scale bar = 30µm. (**D**) Partial view of the gut of *eGFP-cbx1b* KI stage 40 medaka embryo. Gut microvilli are positive for eGFP-Cbx1b (Yellow arrowhead). >10 embryos. Scale bar = 30µm.

**Supplementary Figure 4:**
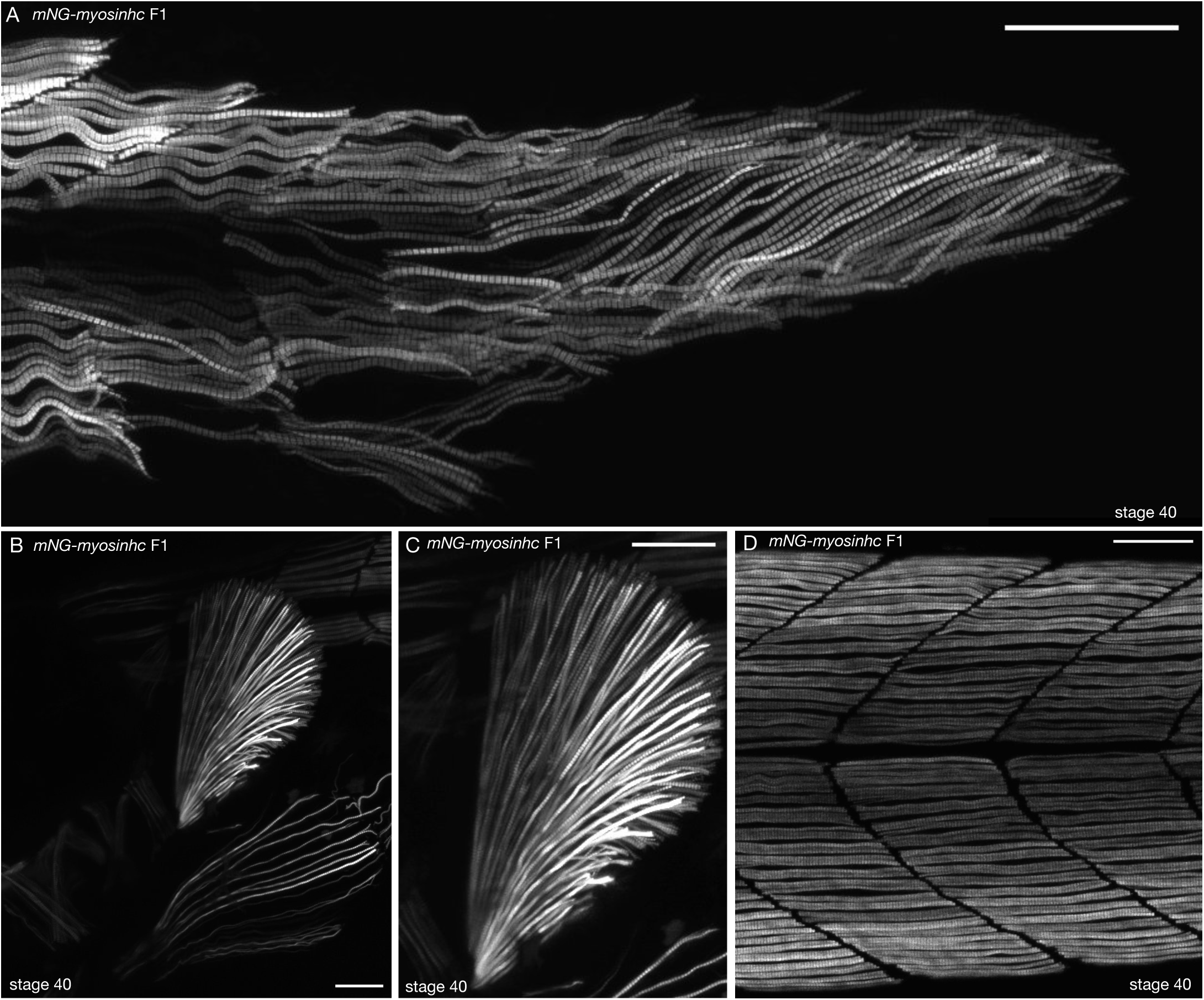
Muscle specific expression of *mNG-myosinhc*. (**A**) Muscle cells of *mNG-myosinhc* F1 stage 40 medaka embryo. mNG-Myosinhc is expressed in muscle cells of the posterior tail region and forms myofibrils, view of myofibrils from Figure 2D. n=10 embryos. Scale bar = 50µm. (**B-C**) Pectoral fin of *mNG-myosinhc* F1 stage 40 medaka embryo. mNG-Myosinhc is expressed in muscles of the pectoral fin, and myofibrils radiate from the base of the pectoral fin. (C) is a close up of (B). n=10 embryos. Scale bar=50µm. (**D**) Body trunk of *mNG-myosinhc* F1 stage 40 medaka embryo. mNG-Myosinhc is expressed in muscle cells in the body trunk. Close-up of muscle cells from Figure 2D. n=10 embryos. Scale bar =50µm

**Supplementary Figure 5:**
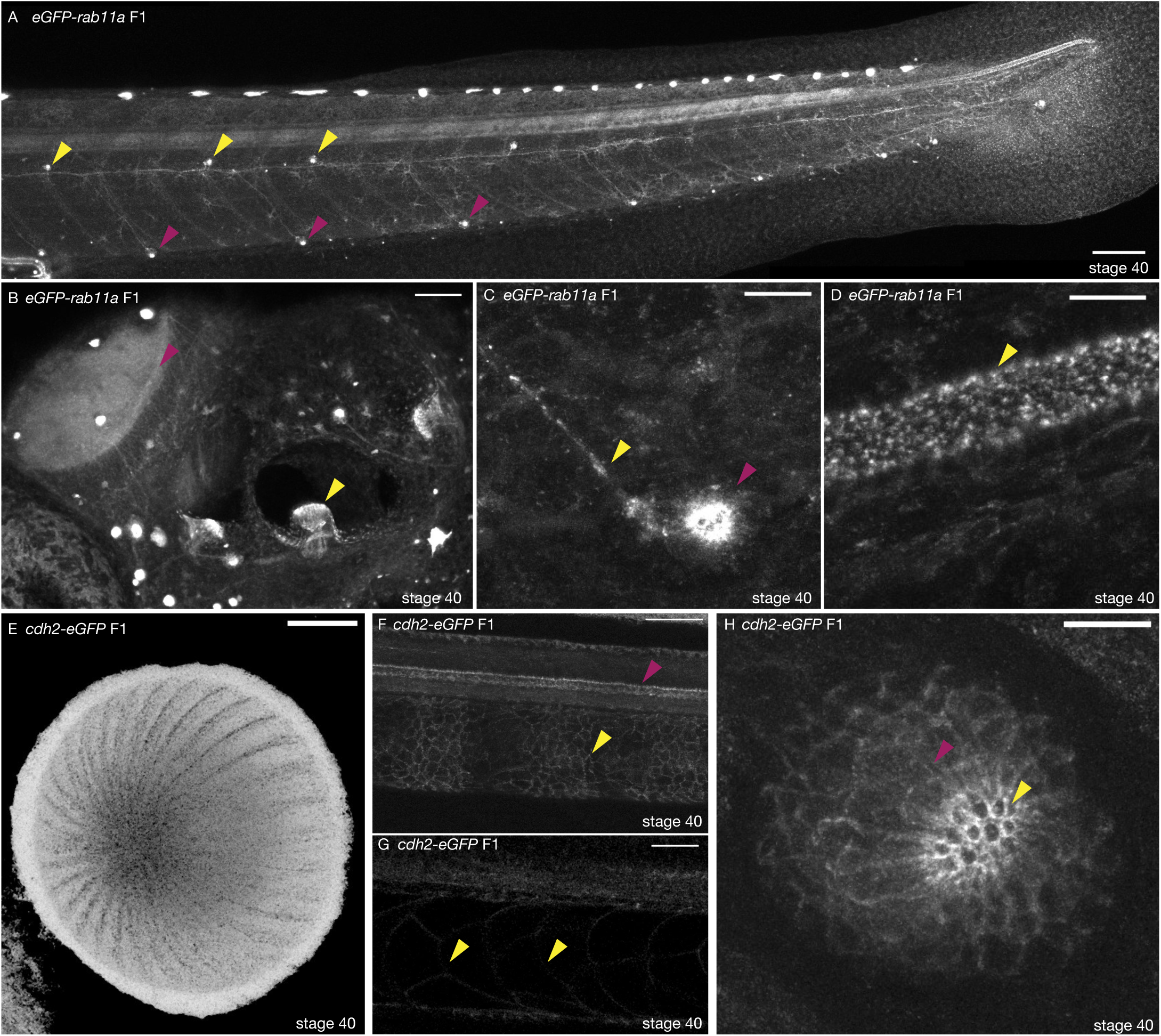
Tissue-specific expression of *eGFP-rab11a* and *cdh2-eGFP* KI lines. (**A**) Neuromasts of *eGFP-rab11a* F1 stage 40 medaka embryo. eGFP-rab11a is strongly expressed in the posterior lateral line neuromasts and posterior lateral line nerve. Ventral primary neuromasts strongly express eGFP-Rab11a (magenta arrowheads), as well as secondary neuromasts located at the horizontal myoseptum (yellow arrowheads). n=10 embryos. Scale bar = 100µm. (**B**) Anterior region of *eGFP-rab11a* F1 stage 40 medaka embryo. eGFP-Rab11a is strongly expressed in the optic tectum (magenta arrowhead) and in the optic vesicle sensory organs (yellow arrowhead). n=10 embryos. Scale bar = 50µm. (**C**) Close-up on a ventral primary neuromast of *eGFP-rab11a* F1 stage 40 medaka embryo. Strong expression of eGFP-Rab11a in neuromast cells (magenta arrowhead) and the lateral line nerve (yellow arrowhead). n=10 embryos. Scale bar = 10µm. (**D**) Spinal cord of *eGFP-rab11a* F1 stage 40 medaka embryo. eGFP-Rab11a is strongly expressed in the spinal cord. n=10 embryos. Scale bar = 10µm. (**E**) Eye of *cdh2-eGFP* F1 stage 40 medaka embryo. Cdh2-eGFP is strongly expressed in elongated cells covering the retina. n=10 embryos. Scale bar = 30µm (**F**) expression of *cdh2-eGFP* F1 in stage 40 medaka embryo. Cdh2-eGFP is expressed in the spinal cord (magenta arrowhead) and notochord sheath cells (yellow arrowhead). n=10 embryos. Scale bar = 50µm. (**G**) Notochord of *cdh2-eGFP* F1 stage 40 medaka embryo. Cdh2-eGFP is expressed in notochord vacuolated cells (yellow arrowhead). n=10 embryos.Scale bar = 30µm. (**H**) Neuromast support and hair cells of *cdh2-eGFP* F1 stage 40 medaka embryo. Neuromast support and hair cells within a mature primary neuromast organ express high levels of Cdh2-eGFP. n=10 embryos. Scale bar =10µm.

**Supplementary Figure 6:**
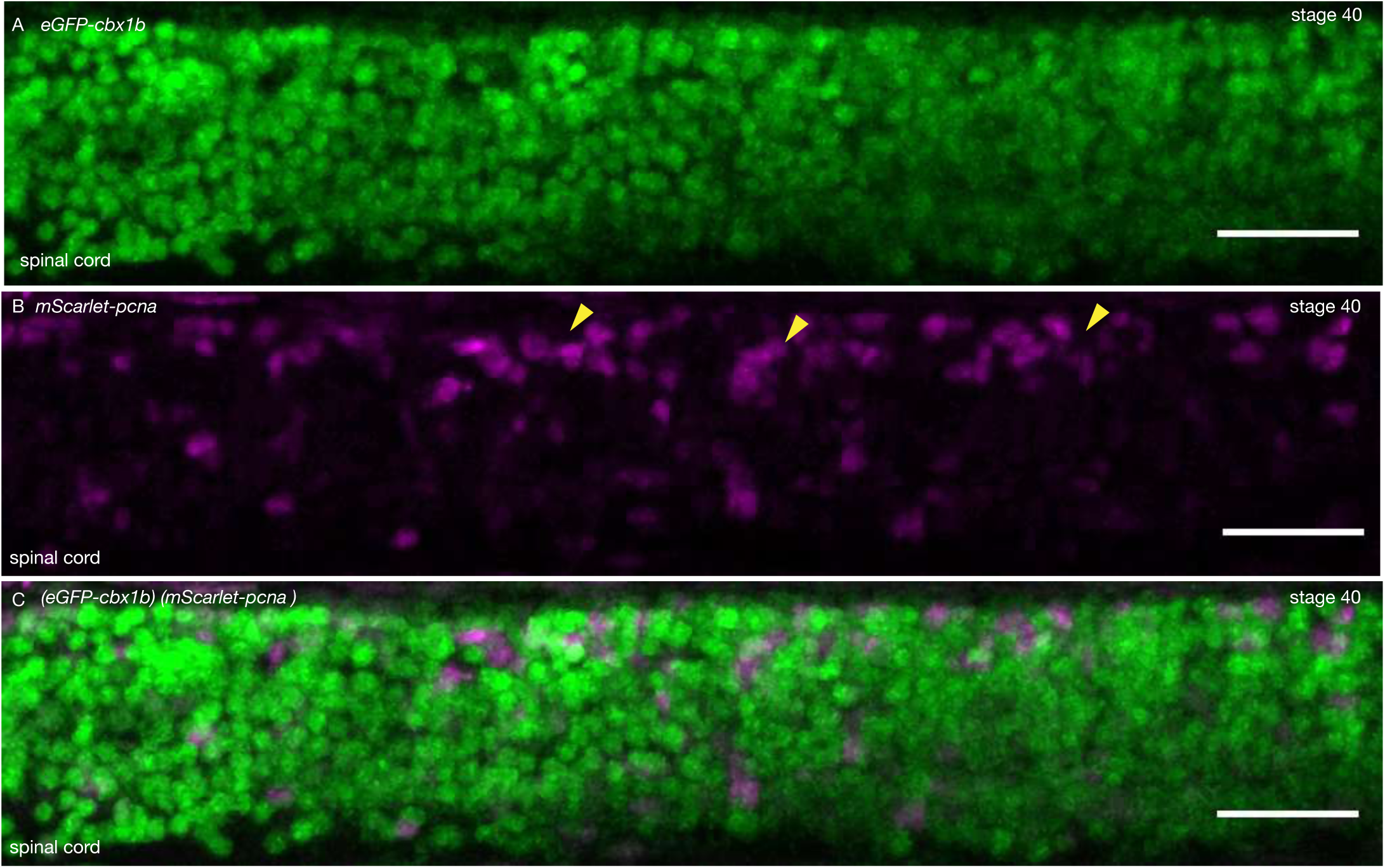
Proliferative cells in the anterior spinal cord. (**A**) Maximum projection of the anterior spinal cord of an (*eGFP-cbx1b*) (*mScarlet-pcna*) double KI stage 40 medaka embryo. eGFP-Cbx1b (Green) is expressed in all anterior spinal cord nuclei. n=4 embryos. Scale bar= 30µm. (**B**) A subset of anterior spinal cord cells are positive for mScarlet-Pcna (Magenta) indicating the existence of cycling cells. mScarlet-Pcna positive cells (yellow arrowheads) are more clustered towards the dorsal side of the spinal cord. n=4 embryos. Scale bar= 30µm. (**C**) Merged image of anterior spinal cord with eGFP-Cbx1b (Green) (A) and mScarlet-Pcna (Magenta) (B). n=4 embryos. Scale bar= 30µm.

**Supplementary Figure 7:**
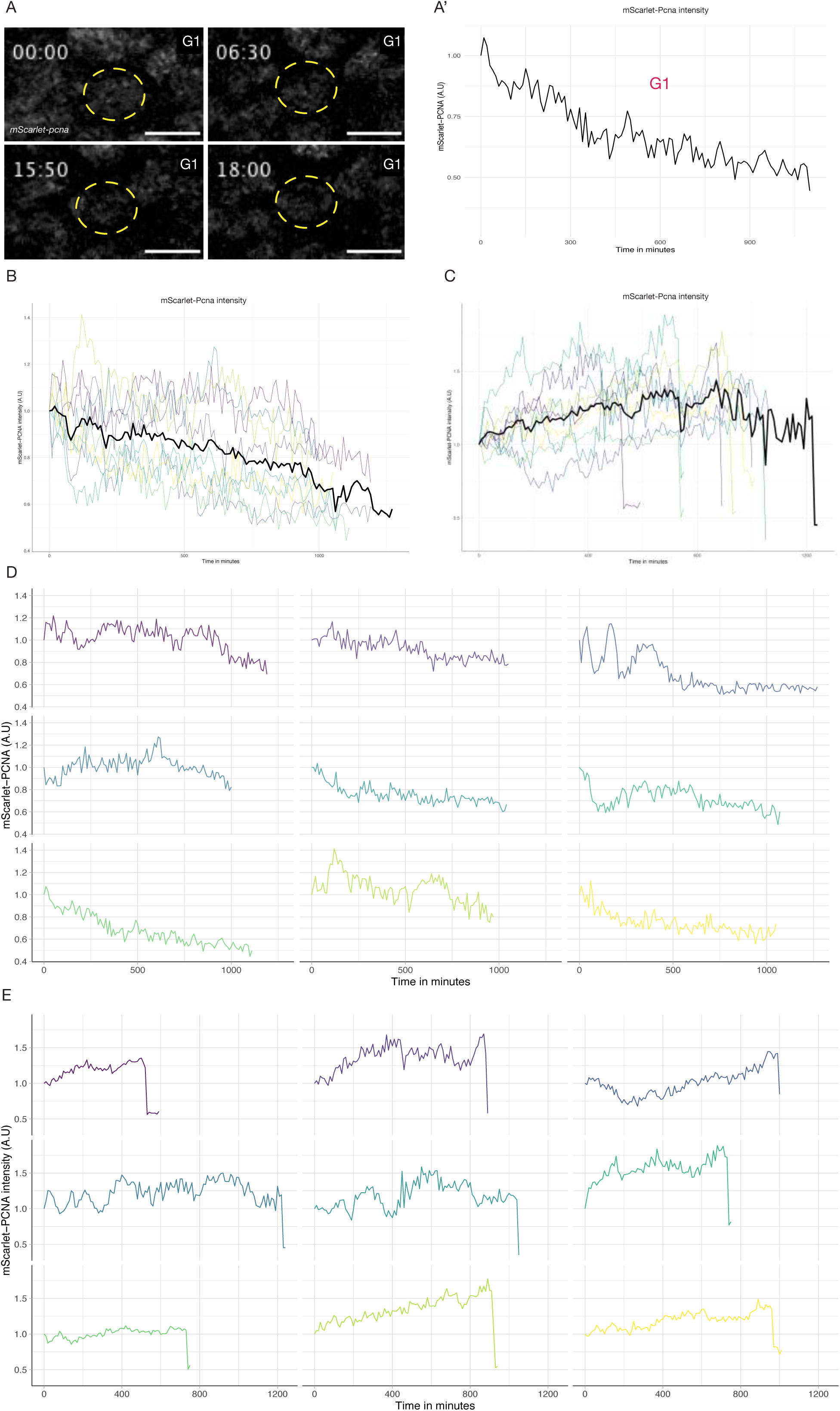
Quantification of endogenous *mScarlet-pcna* levels utilized for cell cycle phase classification. (**A-A’**) Selected frames from a time-lapse of a mScarlet-Pcna positive epithelial cell nucleus that did not undergo cell division over the course of the time-lapse (A, yellow dashed circle). Time in hours. Quantification of normalized mScarlet-Pcna levels shows a decrease of endogenous expression over time indicative of cells in the G1 phase of the cell cycle (A’, n= 9 cells). (**B**) Tracking of endogenous mScarlet-Pcna dynamics in 9 epithelial cell nuclei that do not undergo cell division over the course of the time-lapse (black line = mean). Endogenous mScarlet-Pcna levels decrease over time indicative of cells in the G1 phase of the cell cycle. (**C**) Tracking of endogenous mScarlet-Pcna dynamics in 9 epithelial cell nuclei that undergo cellular division (black line = mean). Endogenous mScarlet-Pcna levels increase over time, indicative of cells in the S phase of the cell cycle. (**D**) Individual traces of normalized mScarlet-Pcna expression in 9 epithelial cells shown in (B). The tracked cells did not divide over the course of the time-lapse. Normalized intensity of mScarlet-Pcna decreases over time indicative of cells in the G1 phase of the cell cycle. (**E**) Individual traces of normalized mScarlet-Pcna expression in 9 epithelial cells shown in (C). The tracked cells underwent a cell division over the course of the time-lapse. Normalized intensity of mScarlet-Pcna increases over time indicative of cells in the S phase of the cell cycle. The beginning of the sharp drop in endogenous mScarlet-Pcna intensity marks entry into M phase.

**Supplementary Figure 8:**
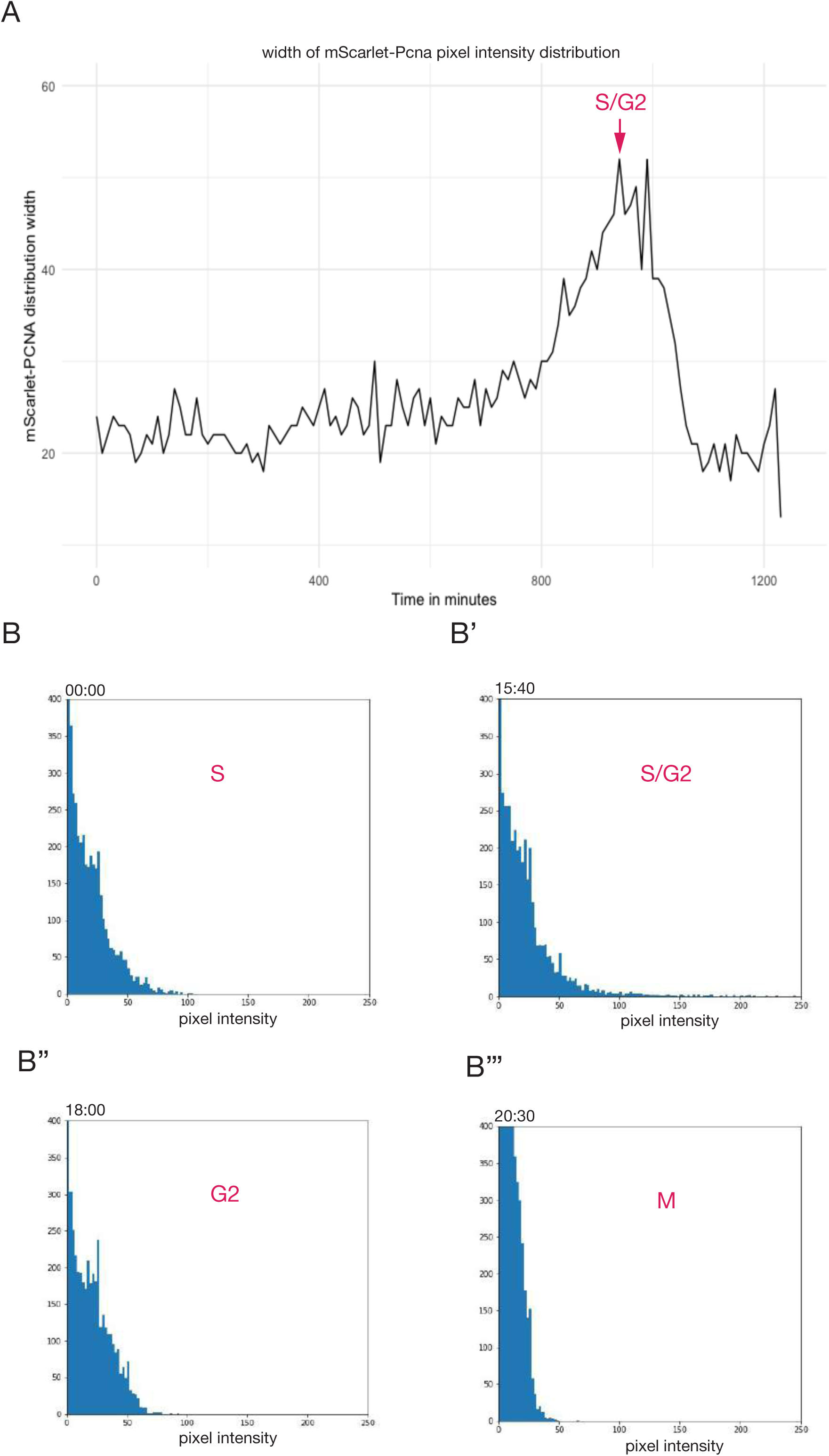
*mScarlet-pcna* histograms of pixel intensity distribution during cell cycle progression. (**A**) Width of mScarlet-Pnca pixel intensity over the course of one cell cycle, the S/G2 transition corresponds to the point of peak pixel intensity distribution within the nucleus and is marked by a red arrow. Data from (A-B) is shown in Figure 4. (**B-B’’’**) Histograms of pixel intensity distribution within the nucleus of the same cell shown in Figure 4A over the course of one cell cycle. The different phases of the cell cycle are marked in red based on the combination of endogenous intensity profiles, 3D surface plots and histograms of pixel intensity distributions. The S/G2 transition is reached in 15:40 and marks the point of peak pixel intensity distribution within the nucleus. M phase is marked by a sharp drop in endogenous mScarlet-Pcna expression. n= 9 epithelial cells.

**Supplementary Movie 1:** Z-stack through the caudal fin region of a stage 39-40 *cbx1-eGFP* medaka embryo. eGFP-Cbx1b is expressed in all nuclei of the different cell types in the caudal fin region. n>10 embryos. Scale bar = 30µm

**Supplementary Movie 2:** Live-imaging in the caudal fin region of a stage 39-40 *eGFP-rab11a* medaka embryo. eGFP-Rab11a is an intra-cellular trafficking marker and localises to intra-cellular vesicles. Notice the dynamics of vesicle trafficking in epithelial cells, neuromasts (magenta arrowhead) and peripheral lateral line nerve (yellow arrowhead). Time in minutes. n=4 embryos. Scale bar =10 µm.

**Supplementary Movie 3:** Live-imaging of skin epithelial cells in the mid-trunk region of a stage 39-40 *eGFP-rab11a* medaka embryo. On the left panel is a merged view of epithelial cells in bright-field and eGFP-Rab11a in green. On the right panel, eGFP-Rab11a in grey scale. eGFP-Rab11a vesicles are prominently displayed as granules within the cytoplasm of epithelial cells. Time in minutes n=4 embryos. Scale bar =10µm.

**Supplementary Movie 4:** Z-stack through the caudal fin region of a stage 39-40 *eGFP-rab11a* medaka embryo. eGFP-Rab11a is strongly expressed in the caudal neuromast and peripheral lateral line nerve. And is also expressed in epithelial cells, the notochord, the spinal cord. n=4 embryos. Scale bar = 30µm.

**Supplementary Movie 5:** Live-imaging of stage 34-35 *g3bp1-eGFP* under normal conditions (temperature 21°C) reveals the cytoplasmic localisation of G3bp1-eGFP in epithelial and muscle cells in the mid trunk region of medaka embryos. Upon stress conditions (temperature shift to 34 °C, 60 minutes after the beginning of the time-lapse), G3bp1-eGFP localization begins to shift into localized clusters of Stress Granule puncta. Time in hours. n=7 embryos. Scale bar = 50µm.

**Supplementary Movie 6:** Live-imaging of stage 34 *mNG-myosinhc* medaka embryo during muscle formation. Muscle cell growth is driven by local buckling of individual muscle cells. Muscle growth and expression of mNG-Myosinhc does not seem to be polarised in an anterior-posterior or dorsal-ventral axis. Instead muscle cells have a heterogenous expression of mNG-Myosinhc that increases as muscle cells grow in length and mature. Time in hours. n=9 embryos .Scale bar =50µm.

**Supplementary Movie 7:** Z-stack through the posterior trunk region of a stage 39-40 *cdh2-eGFP* medaka embryo. Cdh2-eGFP is expressed on the cellular membranes of neuronal tissue including neuromasts, the spinal cord and the notochord. n=3 embryos. Scale bar =50 µm.

**Supplementary Movie 8:** Live-imaging of the dorsal side of a *cdh2-eGFP* medaka embryo at the 12 somite stage reveals the endogenous dynamics of Cdh2-eGFP at high temporal resolution. Time in minutes. n=2. Scale bar=30 µm.

**Supplementary Movie 9:** Live-imaging of stage 39-40 mScarlet-Pcna positive non-dividing epithelial cell nucleus (shown in Figure 4C). Intensity profiles were extracted from within the nucleus (yellow circle). The cell does not divide over the course of the time-lapse. A decrease of mScarlet-Pcna levels over time is indicative of cells in the G1 phase of the cell cycle. Time in hours. n=9 mScarlet-Pcna positive non-dividing epithelial cells. Scale bar =5 µm.

**Supplementary Movie 10:** Live-imaging of stage 39-40 mScarlet-Pcna positive epithelial cell nucleus undergoing cell division (shown in Figure 4D). Intensity profiles were extracted from within the nucleus (yellow circle). An increase of mScarlet-Pcna levels over time is indicative of cells in the S phase of the cell cycle. The appearance of nuclear speckles is indicative of cells in late S phase. The S/G2 transition is marked as the point of peak pixel intensity distribution within the nucleus (yellow circle). The sharp drop of endogenous mScarlet-Pcna levels is indicative of cells in M phase. Time in hours. n=9 mScarlet-Pcna positive and dividing epithelial cells. Scale bar =15 µm.

**Supplementary Movie 11:** Left panel: mScarlet-Pcna positive epithelial cell nucleus undergoing cell division (shown in Figure 4A). Middle panel: histogram of pixel intensity distribution of mScarlet-Pcna within the nucleus of the tracked cell. Right panel: frequency distribution width obtained from the histogram of pixel intensity distribution. Peak pixel intensity distribution is reached at 15:40h and is used to mark the S/G2 transition. Time in hours. n=9 dividing epithelial cells. Scale bar =15 µm.

**Supplementary Movie 12:** Left panel: 3D surface plot of mScarlet-Pcna positive epithelial cell nucleus undergoing cell division (from Figure 4A). Middle panel: normalized endogenous mScarlet-Pcna levels of epithelial cell (from Figure 4A). Right panel: histogram of pixel intensity distribution. Time in hours. n=9 epithelial cells that undergo cell division.

**Supplementary File S1:** Detailed protocol for cloning-free CRISPR insertion of fluorescent reporters in medaka

**Supplementary File S2:** Detailed sequence design for *mNeonGreen-HAtag-Linker-myosinhc* tagging

**Table 1:** Quantification of targeting efficiency

**Table S1:** Experimental and targeted *loci* details

**Table S2:** Donor templates used in this study

**Table S3:** Primers used in this study

**Table S4:** Plasmids used in this study

**Table S5:** sgRNA used in this study

**Table S6:** Medaka lines generated and maintained in this study

## Materials and Methods

### Animal husbandry and ethics

Medaka (*Oryzias latipes*) (Iwamatsu, 2004, Naruse et al., 2004, Kasahara et al., 2007) were maintained as closed stocks in a fish facility built according to the European Union animal welfare standards and all animal experiments were performed in accordance with European Union animal welfare guidelines. Animal experimentation was approved by The EMBL Institutional Animal Care and Use Committee (IACUC) project code: 20/001_HD_AA. Fishes were maintained in a constant recirculating system at 27-28°C with a 14hr light / 10hr dark cycle.

### Cloning-free CRISPR/Cas9 Knock-Ins

A detailed step-by-step protocol for the cloning-free approach is provided in Files S1/S2. A detailed list of all repair donors, PCR primers, fluorescent protein sequences and sgRNAs used is provided in Tables S1-S6. Briefly, for the preparation of Cas9-mSA mRNA: the pCS2+Cas9-mSA plasmid was a gift from Janet Rossant (Addgene #103882) (Gu et al., 2018). 6-8 µg of Cas9-mSA plasmid was linearized by Not1-HF restriction enzyme (NEB #R3189S). The 8.8kb linearized fragment was cut out from a 1.5% agarose gel and DNA was extracted using QIAquick Gel Extraction Kit (Qiagen #28115). *In vitro* transcription was performed using mMachine SP6 Transcription Kit (Invitrogen #AM1340) following the manufacturer’s guidelines. RNA cleanup was performed using RNAeasy Mini Kit (Qiagen #74104). sgRNAs were manually selected using previously published recommendations (Paix et al., 2017a, Paix et al., 2019, Doench et al., 2016, Gagnon et al., 2014) and *in silico* validated using CCTop and CHOPCHOP (Labun et al., 2019, Stemmer et al., 2015) (Table S5). The genomic coordinates of all genes targeted can be found in Table S1. Synthetic sgRNAs used in this study were ordered from Sigma-Aldrich (spyCas9 sgRNA, 3nmole, HPLC purification, no modification). PCR repair donor fragments were prepared as described previously (Paix et al., 2014, Paix et al., 2017b, Paix et al., 2015, Paix et al., 2016) and a detailed protocol is provided in File S1. Briefly the design includes approx. 30-40bp of homology arms and a fluorescent protein sequence with no ATG or Stop codon (Tables S2/S4). PCR amplifications were performed using Phusion or Q5 high fidelity DNA polymerase (NEB Phusion Master Mix with HF buffer #M0531L or NEB Q5 Master Mix # M0492L). MinElute PCR Purification Kit (Qiagen #28004) was used for PCR purification. Primers were ordered from Sigma-Aldrich (25nmole scale, desalted) and contained Biotin moiety on the 5’ ends for repair donor synthesis. A list of all primers and fluorescent protein sequences used in this study can be found in Table S3/S4. The injection mix in medaka contains the sgRNA (15-20 ng/ul) + Cas9-mSA mRNA (150 ng/ul) + repair donor template (8-10 ng/ul). For injections, male and female medakas are added to the same tank and fertilized eggs collected 20 minutes later. The mix is injected in 1-cell staged medaka embryos (Iwamatsu, 2004), and embryos are raised at 28°C in 1XERM (Seleit et al., 2017a, Seleit et al., 2017b, Rembold et al., 2006). A list of KI lines generated and maintained in this study can be found in Table S6.

### Live-imaging sample preparation

Embryos were prepared for live-imaging as previously described (Seleit et al., 2017a, Seleit et al., 2017b). 1X Tricaine (Sigma-Aldrich #A5040-25G) was used to anesthetize dechorionated medaka embryos (20 mg/ml – 20X stock solution diluted in 1XERM). Anesthetized embryos were then mounted in low melting agarose (0.6 to 1%) (Biozyme Plaque Agarose #840101). Imaging was done on glass-bottomed dishes (MatTek Corporation Ashland, MA 01721, USA). For *g3bp1-eGFP* live-imaging, temperature was changed from 21°C to 34°C after one hour of imaging.

### Microscopy and data analysis

For all embryo screening, a Nikon SMZ18 fluorescence stereoscope was used. All live-imaging, except for *g3bp1-eGFP* and *cdh2-eGFP embryos,* was done on a laser-scanning confocal Leica SP8 (CSU, White Laser) microscope, 20x and 40x objectives were used during image acquisition depending on the experimental sample. For the SP8 confocal equipped with a white laser, the laser emission was matched to the spectral properties of the fluorescent protein of interest. *g3bp1-eGFP* line live-imaging was performed using a Zeiss LSM780 laser-scanning confocal with a temperature control box and an Argon laser at 488 nm, imaged through a 20x plan apo objective (numerical aperture 0.8). For *cdh2-eGFP* 4D live-imaging was performed on a Luxendo TruLive SPIM system using a 30X objective. Open-source standard ImageJ/Fiji software (Schindelin et al., 2012) was used for analysis and editing of all images post image acquisition. Stitching was performed using standard 2D and 3D stitching plug-ins on ImageJ/Fiji. For quantitative values on endogenous *mScarlet-pcna* dynamics ROI manager in ImageJ/Fiji was used to define fluorescence intensity within the nucleus of tracked cells (Yellow circle in Figure 4 and Supplementary Movie S10/11), fluorescent intensity measurements were then extracted from the time-series and the data was normalized by dividing on the initial intensity value in each time-lapse movies. Data was plotted using R software. Pixel intensity distribution within nuclei were analyzed using a custom python based script. Individual live-cell tracks were plotted using PlotTwist (Goedhart, 2020).

### Fin-clips, genotyping and sanger sequencing

Individual adult F1 fishes were fin-clipped for genotyping PCRs. Briefly, fish were anesthetized in 1X Tricaine solution. A small part of the caudal fin was cut by sharp scissors and placed in a 2ml Eppendorf tube containing 50ul of fin-clip buffer. The fish were recovered in small beakers and were transferred back to their tanks. Eppendorf tubes were then incubated overnight at 65°C. 100µl of H_2_O was then added to each tube and then the tubes were incubated for 10-15 min at 90°C. Tubes were then centrifuged for 30 minutes at 10,000 rpm in a standard micro-centrifuge. Supernatant was used for subsequent PCRs. Fin-clip buffer is composed of 100 ml 2M Tris pH 8.0, 5 ml 0.5M EDTA pH 8.0, 15 ml 5M NaCl, 2.5 ml 20% SDS, H2O to 500 ml, sterile filtered. 50 ul of proteinase K (20 mg/ml) was added to 1 ml fin clip buffer before use. 2ul of genomic DNA from fin-clips was used for genotyping PCRs. A list of all genotyping primers used in this study can be found in Table S3. After PCRs the edited and WT amplicons were sent to Sanger sequencing (Eurofins Genomics). Sequences were analyzed using Geneious software (Figure S2). In-frame integrations were confirmed by sequencing for *eGFP-cbx1b, mScarlet-pcna, mNeonGreen-myosinhc* and *eGFP-rab11a.* We were able to detect an internal partial duplication of the 5’ homology arm in the *mScarlet-pcna* line that does not affect the protein coding sequence nor the 5’ extremity of the homology arm itself. Specifically, 22 base pairs upstream of the Start codon of *pcna* (and within the 5’ homology arm); we detect a 21 bp partial duplication of the 5’ homology arm and a 7bp insertion GGTCGAC indicative that the repair mechanism involved can lead to errors (Paix et al., 2017a). The 5’ homology junction itself is unaltered and precise.

### Whole Genome Sequencing (WGS)

5 to 10 positive F1 medaka embryos (originating from the same F0 founder) of the *eGFP-cbx1b, mScarlet-pcna* and *mNeonGreen-myosinhc* lines were snap frozen in liquid nitrogen and kept at -80°C in 1.5ml Eppendorf tubes. Genomic DNA was extracted using DNeasy Blood and Tissue Kit (Qiagen #69504) according to the manufacturer’s guidelines. The libraries were prepared on a liquid handling system (Beckman i7 series) using 200 ng of sheared gDNA and 10 PCR cycles using the NEBNext Ultra II DNA Library Prep Kit for Illumina (NEB #E7645S). The DNA libraries were indexed with unique dual barcodes (8bp long), pooled together and then sequenced using an Illumina NextSeq550 instrument with a 150 PE mid-mode in paired-end mode with a read length of 150bp. Sequenced reads were aligned to the *Oryzias latipes* reference genome (*Ensembl!* Assembly version ASM223467v1) using BWA mem version 0.7.17 with default settings (Li and Durbin, 2009). The reference genome was augmented with the known inserts for *eGFP*, *mScarlet* and *mNeonGreen* to facilitate a direct integration discovery using standard inter-chromosomal structural variant predictions. The insert sequences are provided in Tables S2/S4. After the genome alignment, reads were sorted and indexed using SAMtools (Li et al., 2009). Quality control and coverage analyses were performed using the Alfred qc subcommand (Rausch et al., 2019). For Structural Variant (SV) discovery, aligned reads were processed with DELLY v0.8.7 (Rausch et al., 2012) using paired-end mapping and split-read analysis. SVs were filtered for inter-chromosomal SVs with one breakpoint in one of the additional insert sequences (*eGFP*, *mScarlet* and *mNeonGreen*). Plots shown in Figure S2 are adapted from Integrative Genomics Viewer (IGV) (Thorvaldsdottir et al., 2013). The estimated genomic coordinates for integration are: *eGFP-cbx1b* (chr19:19,074,552), *mScarlet-pcna* (chr9:6,554,003) and *mNeonGreen-myosinhc* (chr8:8,975,799). Coverage of *eGFP-cbx1b*_gDNA1 is 20.4X and *eGFP-cbx1b*_gDNA2 is 23.6X. Coverage of *mScarlet-pcna* is 14.4X. Coverage of *mNeonGreen-myosinhc* is 14.5X. Raw sequencing data was deposited in European Nucleotide Archive (ENA) under study number ERP127162. Accession numbers are: *eGFP-cbx1b*(1) ERS5796960 (SAMEA8109891), *eGFP-cbx1b*(2) ERS5796961 (SAMEA8109892), *mScarlet-pcna* ERS5796962 (SAMEA8109893) and *mNeonGreen-myosinhc* ERS5796963 (SAMEA8109894).

## Acknowledgments

We would like to thank all members of the Aulehla lab for the fruitful discussions on the work presented here. We would like to thank Aissam Ikmi for input on the manuscript. We would like to thank Takehito Tomita for help with python scripts. The European Molecular Biology Laboratory (EMBL) Heidelberg Genecore is acknowledged for support in WGS data acquisition and analysis. We would like to thank Vladimir Benes and members of his team at Genecore EMBL Heidelberg for continuous help and support, Tobias Rausch for computational work on the WGS data and Mireia Osuna Lopez for help in library preparation of WGS DNA. In addition, we would like to thank all animal-care takers at EMBL Heidelberg and in particular Sabine Goergens for excellent support. We would also like to thank Addgene for access to plasmids. This work was supported by: the European Molecular Biology Laboratory (EMBL), Heidelberg and the EMBL interdisciplinary Postdoc (EIPOD4) under Marie Sklodowska-Curie Actions Cofund (grant agreement number 847543) fellowship for funding to Ali Seleit. This work also received support from the European Research Council under an ERC consolidator grant agreement n.866537 to A.A.

## References

Adams, D. S., Keller, R. & Koehl, M. A. 1990. The mechanics of notochord elongation, straightening and stiffening in the embryo of Xenopus laevis. Development, 110, 115–30.

Akitake, C. M., Macurak, M., Halpern, M. E. & Goll, M. G. 2011. Transgenerational analysis of transcriptional silencing in zebrafish. Dev Biol, 352, 191–201.

Alunni, A., Hermel, J. M., Heuze, A., Bourrat, F., Jamen, F. & Joly, J. S. 2010. Evidence for neural stem cells in the medaka optic tectum proliferation zones. Dev Neurobiol, 70, 693–713.

Araujo, A. R., Gelens, L., Sheriff, R. S. & Santos, S. D. 2016. Positive Feedback Keeps Duration of Mitosis Temporally Insulated from Upstream Cell-Cycle Events. Mol Cell, 64, 362–375.

Auer, T. O. & Del Bene, F. 2014. Crispr/Cas9 and Talen-mediated knock-in approaches in zebrafish. Methods, 69, 142–50.

Auer, T. O., Duroure, K., De Cian, A., Concordet, J. P. & Del Bene, F. 2014. Highly efficient Crispr/Cas9-mediated knock-in in zebrafish by homology-independent Dna repair. Genome Res, 24, 142–53.

Bajar, B. T., Lam, A. J., Badiee, R. K., Oh, Y. H., Chu, J., Zhou, X. X., Kim, N., Kim, B. B., Chung, M., Yablonovitch, A. L., Cruz, B. F., Kulalert, K., Tao, J. J., Meyer, T., Su, X. D. & Lin, M. Z. 2016. Fluorescent indicators for simultaneous reporting of all four cell cycle phases. Nat Methods, 13, 993–996.

Barr, A. R., Heldt, F. S., Zhang, T., Bakal, C. & Novak, B. 2016. A Dynamical Framework for the All-or-None G1/S Transition. Cell Syst, 2, 27–37.

Bindels, D. S., Haarbosch, L., Van Weeren, L., Postma, M., Wiese, K. E., Mastop, M., Aumonier, S., Gotthard, G., Royant, A., Hink, M. A. & Gadella, T. W., Jr. 2017. mScarlet: a bright monomeric red fluorescent protein for cellular imaging. Nat Methods, 14, 53–56.

Buczacki, S. J., Zecchini, H. I., Nicholson, A. M., Russell, R., Vermeulen, L., Kemp, R. & Winton, D. J. 2013. Intestinal label-retaining cells are secretory precursors expressing Lgr5. Nature, 495, 65–9.

Burket, C. T., Montgomery, J. E., Thummel, R., Kassen, S. C., Lafave, M. C., Langenau, D. M., Zon, L. I. & Hyde, D. R. 2008. Generation and characterization of transgenic zebrafish lines using different ubiquitous promoters. Transgenic Res, 17, 265–79.

Buttitta, L. A., Katzaroff, A. J. & Edgar, B. A. 2010. A robust cell cycle control mechanism limits E2F-induced proliferation of terminally differentiated cells in vivo. J Cell Biol, 189, 981–96.

Ceccaldi, R., Rondinelli, B. & D’andrea, A. D. 2016. Repair Pathway Choices and Consequences at the Double-Strand Break. Trends Cell Biol, 26, 52–64.

Centanin, L., Ander, J. J., Hoeckendorf, B., Lust, K., Kellner, T., Kraemer, I., Urbany, C., Hasel, E., Harris, W. A., Simons, B. D. & Wittbrodt, J. 2014. Exclusive multipotency and preferential asymmetric divisions in post-embryonic neural stem cells of the fish retina. Development, 141, 3472–82.

Chopra, A., Tabdanov, E., Patel, H., Janmey, P. A. & Kresh, J. Y. 2011. Cardiac myocyte remodeling mediated by N-cadherin-dependent mechanosensing. Am J Physiol Heart Circ Physiol, 300, H1252–66.

Chudakov, D. M., Matz, M. V., Lukyanov, S. & Lukyanov, K. A. 2010. Fluorescent proteins and their applications in imaging living cells and tissues. Physiol Rev, 90, 1103–63.

Cong, L., Ran, F. A., Cox, D., Lin, S., Barretto, R., Habib, N., Hsu, P. D., Wu, X., Jiang, W., Marraffini, L. A. & Zhang, F. 2013. Multiplex genome engineering using Crispr/Cas systems. Science, 339, 819–23.

Cristea, S., Freyvert, Y., Santiago, Y., Holmes, M. C., Urnov, F. D., Gregory, P. D. & Cost, G. J. 2013. In vivo cleavage of transgene donors promotes nuclease-mediated targeted integration. Biotechnol Bioeng, 110, 871–80.

Cullen, P. J. & Steinberg, F. 2018. To degrade or not to degrade: mechanisms and significance of endocytic recycling. Nat Rev Mol Cell Biol, 19, 679–696.

Danner, E., Bashir, S., Yumlu, S., Wurst, W., Wefers, B. & Kuhn, R. 2017. Control of gene editing by manipulation of Dna repair mechanisms. Mamm Genome, 28, 262–274.

Decker, C. J. & Parker, R. 2012. P-bodies and stress granules: possible roles in the control of translation and mRNA degradation. Cold Spring Harb Perspect Biol, 4, a012286.

Desclozeaux, M., Venturato, J., Wylie, F. G., Kay, J. G., Joseph, S. R., Le, H. T. & Stow, J. L. 2008. Active Rab11 and functional recycling endosome are required for E-cadherin trafficking and lumen formation during epithelial morphogenesis. Am J Physiol Cell Physiol, 295, C545–56.

Dickinson, D. J., Pani, A. M., Heppert, J. K., Higgins, C. D. & Goldstein, B. 2015. Streamlined Genome Engineering with a Self-Excising Drug Selection Cassette. Genetics, 200, 1035–49.

Doench, J. G., Fusi, N., Sullender, M., Hegde, M., Vaimberg, E. W., Donovan, K. F., Smith, I., Tothova, Z., Wilen, C., Orchard, R., Virgin, H. W., Listgarten, J. & Root, D. E. 2016. Optimized sgRNA design to maximize activity and minimize off-target effects of Crispr-Cas9. Nat Biotechnol, 34, 184–191.

Dolfi, L., Ripa, R., Antebi, A., Valenzano, D. R. & Cellerino, A. 2019. Cell cycle dynamics during diapause entry and exit in an annual killifish revealed by Fucci technology. Evodevo, 10, 29.

Dufourcq, P., Roussigne, M., Blader, P., Rosa, F., Peyrieras, N. & Vriz, S. 2006. Mechano-sensory organ regeneration in adults: the zebrafish lateral line as a model. Mol Cell Neurosci, 33, 180–7.

Fu, Y., Foden, J. A., Khayter, C., Maeder, M. L., Reyon, D., Joung, J. K. & Sander, J. D. 2013. High-frequency off-target mutagenesis induced by Crispr-Cas nucleases in human cells. Nat Biotechnol, 31, 822–6.

Gagnon, J. A., Valen, E., Thyme, S. B., Huang, P., Akhmetova, L., Pauli, A., Montague, T. G., Zimmerman, S., Richter, C. & Schier, A. F. 2014. Efficient mutagenesis by Cas9 protein-mediated oligonucleotide insertion and large-scale assessment of single-guide RNAs. PLoS One, 9, e98186.

Garcia, J., Bagwell, J., Njaine, B., Norman, J., Levic, D. S., Wopat, S., Miller, S. E., Liu, X., Locasale, J. W., Stainier, D. Y. R. & Bagnat, M. 2017. Sheath Cell Invasion and Trans-differentiation Repair Mechanical Damage Caused by Loss of Caveolae in the Zebrafish Notochord. Curr Biol, 27, 1982–1989 e3.

Gibson, T. J., Seiler, M. & Veitia, R. A. 2013. The transience of transient overexpression. Nat Methods, 10, 715–21.

Gilmore, J. M., Sardiu, M. E., Groppe, B. D., Thornton, J. L., Liu, X., Dayebgadoh, G., Banks, C. A., Slaughter, B. D., Unruh, J. R., Workman, J. L., Florens, L. & Washburn, M. P. 2016. WDR76 Co-Localizes with Heterochromatin Related Proteins and Rapidly Responds to Dna Damage. PLoS One, 11, e0155492.

Goedhart, J. 2020. PlotTwist: A web app for plotting and annotating continuous data. PLoS Biol, 18, e3000581.

Goll, M. G., Anderson, R., Stainier, D. Y., Spradling, A. C. & Halpern, M. E. 2009. Transcriptional silencing and reactivation in transgenic zebrafish. Genetics, 182, 747–55.

Gratz, S. J., Ukken, F. P., Rubinstein, C. D., Thiede, G., Donohue, L. K., Cummings, A. M. & O’Connor-Giles, K. M. 2014. Highly specific and efficient Crispr/Cas9-catalyzed homology-directed repair in Drosophila. Genetics, 196, 961–71.

Gu, B., Posfai, E. & Rossant, J. 2018. Efficient generation of targeted large insertions by microinjection into two-cell-stage mouse embryos. Nat Biotechnol, 36, 632–637.

Guarino, A. M., Mauro, G. D., Ruggiero, G., Geyer, N., Delicato, A., Foulkes, N. S., Vallone, D. & Calabro, V. 2019. Yb-1 recruitment to stress granules in zebrafish cells reveals a differential adaptive response to stress. Sci Rep, 9, 9059.

Gutierrez-Triana, J. A., Tavhelidse, T., Thumberger, T., Thomas, I., Wittbrodt, B., Kellner, T., Anlas, K., Tsingos, E. & Wittbrodt, J. 2018. Efficient single-copy Hdr by 5’ modified long dsDNA donors. Elife, 7.

Hackett, C. S., Geurts, A. M. & Hackett, P. B. 2007. Predicting preferential Dna vector insertion sites: implications for functional genomics and gene therapy. Genome Biol, 8 Suppl 1, S12.

Halbleib, J. M. & Nelson, W. J. 2006. Cadherins in development: cell adhesion, sorting, and tissue morphogenesis. Genes Dev, 20, 3199–214.

Harrington, M. J., Hong, E., Fasanmi, O. & Brewster, R. 2007. Cadherin-mediated adhesion regulates posterior body formation. Bmc Dev Biol, 7, 130.

Hartman, M. A. & Spudich, J. A. 2012. The myosin superfamily at a glance. J Cell Sci, 125, 1627–32.

Held, M., Schmitz, M. H., Fischer, B., Walter, T., Neumann, B., Olma, M. H., Peter, M., Ellenberg, J. & Gerlich, D. W. 2010. CellCognition: time-resolved phenotype annotation in high-throughput live cell imaging. Nat Methods, 7, 747–54.

Hisano, Y., Sakuma, T., Nakade, S., Ohga, R., Ota, S., Okamoto, H., Yamamoto, T. & Kawahara, A. 2015. Precise in-frame integration of exogenous Dna mediated by Crispr/Cas9 system in zebrafish. Sci Rep, 5, 8841.

Hoshijima, K., Jurynec, M. J. & Grunwald, D. J. 2016. Precise Editing of the Zebrafish Genome Made Simple and Efficient. Dev Cell, 36, 654–67.

Hoshijima, K., Jurynec, M. J., Klatt Shaw, D., Jacobi, A. M., Behlke, M. A. & Grunwald, D. J. 2019. Highly Efficient Crispr-Cas9-Based Methods for Generating Deletion Mutations and F0 Embryos that Lack Gene Function in Zebrafish. Dev Cell, 51, 645–657 e4.

Irvine, K., Stirling, R., Hume, D. & Kennedy, D. 2004. Rasputin, more promiscuous than ever: a review of G3BP. Int J Dev Biol, 48, 1065–77.

Iwamatsu, T. 2004. Stages of normal development in the medaka Oryzias latipes. Mech Dev, 121, 605–18.

Jasin, M. & Haber, J. E. 2016. The democratization of gene editing: Insights from site-specific cleavage and double-strand break repair. Dna Repair (Amst), 44, 6–16.

Jinek, M., Chylinski, K., Fonfara, I., Hauer, M., Doudna, J. A. & Charpentier, E. 2012. A programmable dual-Rna-guided Dna endonuclease in adaptive bacterial immunity. Science, 337, 816–21.

Jones, J. E. & Corwin, J. T. 1993. Replacement of lateral line sensory organs during tail regeneration in salamanders: identification of progenitor cells and analysis of leukocyte activity. J Neurosci, 13, 1022–34.

Kanca, O., Zirin, J., Garcia-Marques, J., Knight, S. M., Yang-Zhou, D., Amador, G., Chung, H., Zuo, Z., Ma, L., He, Y., Lin, W. W., Fang, Y., Ge, M., Yamamoto, S., Schulze, K. L., Hu, Y., Spradling, A. C., Mohr, S. E., Perrimon, N. & Bellen, H. J. 2019. An efficient Crispr-based strategy to insert small and large fragments of Dna using short homology arms. Elife, 8.

Kasahara, M., Naruse, K., Sasaki, S., Nakatani, Y., Qu, W., Ahsan, B., Yamada, T., Nagayasu, Y., Doi, K., Kasai, Y., Jindo, T., Kobayashi, D., Shimada, A., Toyoda, A., Kuroki, Y., Fujiyama, A., Sasaki, T., Shimizu, A., Asakawa, S., Shimizu, N., Hashimoto, S., Yang, J., Lee, Y., Matsushima, K., Sugano, S., Sakaizumi, M., Narita, T., Ohishi, K., Haga, S., Ohta, F., Nomoto, H., Nogata, K., Morishita, T., Endo, T., Shin, I. T., Takeda, H., Morishita, S. & Kohara, Y. 2007. The medaka draft genome and insights into vertebrate genome evolution. Nature, 447, 714–9.

Kim, J. H., Lee, S. R., Li, L. H., Park, H. J., Park, J. H., Lee, K. Y., Kim, M. K., Shin, B. A. & Choi, S. Y. 2011. High cleavage efficiency of a 2A peptide derived from porcine teschovirus-1 in human cell lines, zebrafish and mice. PLoS One, 6, e18556.

Kroll, F., Powell, G. T., Ghosh, M., Gestri, G., Antinucci, P., Hearn, T. J., Tunbak, H., Lim, S., Dennis, H. W., Fernandez, J. M., Whitmore, D., Dreosti, E., Wilson, S. W., Hoffman, E. J. & Rihel, J. 2021. A simple and effective F0 knockout method for rapid screening of behaviour and other complex phenotypes. Elife, 10.

Kuo, C. T., You, G. T., Jian, Y. J., Chen, T. S., Siao, Y. C., Hsu, A. L. & Ching, T. T. 2020. Ampk-mediated formation of stress granules is required for dietary restriction-induced longevity in Caenorhabditis elegans. Aging Cell, 19, e13157.

Labun, K., Montague, T. G., Krause, M., Torres Cleuren, Y. N., Tjeldnes, H. & Valen, E. 2019. Chopchop v3: expanding the Crispr web toolbox beyond genome editing. Nucleic Acids Res, 47, W171–W174.

Lavker, R. M. & Sun, T. T. 2003. Epithelial stem cells: the eye provides a vision. Eye (Lond), 17, 937–42.

Leckband, D. E. & De Rooij, J. 2014. Cadherin adhesion and mechanotransduction. Annu Rev Cell Dev Biol, 30, 291–315.

Leonetti, M. D., Sekine, S., Kamiyama, D., Weissman, J. S. & Huang, B. 2016. A scalable strategy for high-throughput Gfp tagging of endogenous human proteins. Proc Natl Acad Sci U S A, 113, E3501–8.

Leonhardt, H., Rahn, H. P., Weinzierl, P., Sporbert, A., Cremer, T., Zink, D. & Cardoso, M. C. 2000. Dynamics of Dna replication factories in living cells. J Cell Biol, 149, 271–80.

Leung, L., Klopper, A. V., Grill, S. W., Harris, W. A. & Norden, C. 2011. Apical migration of nuclei during G2 is a prerequisite for all nuclear motion in zebrafish neuroepithelia. Development, 138, 5003–13.

Levic, D. S., Yamaguchi, N., Wang, S., Knaut, H. & Bagnat, M. 2021. Knock-in tagging in zebrafish facilitated by insertion into non-coding regions. bioRxiv, 2021.07.08.451679.

Li, H. & Durbin, R. 2009. Fast and accurate short read alignment with Burrows-Wheeler transform. Bioinformatics, 25, 1754–60.

Li, H., Handsaker, B., Wysoker, A., Fennell, T., Ruan, J., Homer, N., Marth, G., Abecasis, G., Durbin, R. & Genome Project Data Processing, S. 2009. The Sequence Alignment/Map format and SAMtools. Bioinformatics, 25, 2078–9.

Li, W., Zhang, Y., Han, B., Li, L., Li, M., Lu, X., Chen, C., Lu, M., Zhang, Y., Jia, X., Zhu, Z., Tong, X. & Zhang, B. 2019. One-step efficient generation of dual-function conditional knockout and geno-tagging alleles in zebrafish. Elife, 8.

Lim, Y. W., Lo, H. P., Ferguson, C., Martel, N., Giacomotto, J., Gomez, G. A., Yap, A. S., Hall, T. E. & Parton, R. G. 2017. Caveolae Protect Notochord Cells against Catastrophic Mechanical Failure during Development. Curr Biol, 27, 1968–1981 e7.

Lisby, M. & Rothstein, R. 2004. Dna damage checkpoint and repair centers. Curr Opin Cell Biol, 16, 328–34.

Loison, O., Weitkunat, M., Kaya-Copur, A., Nascimento Alves, C., Matzat, T., Spletter, M. L., Luschnig, S., Brasselet, S., Lenne, P. F. & Schnorrer, F. 2018. Polarization-resolved microscopy reveals a muscle myosin motor-independent mechanism of molecular actin ordering during sarcomere maturation. PLoS Biol, 16, e2004718.

Lomberk, G., Wallrath, L. & Urrutia, R. 2006. The Heterochromatin Protein 1 family. Genome Biol, 7, 228.

Lu, C. P., Polak, L., Rocha, A. S., Pasolli, H. A., Chen, S. C., Sharma, N., Blanpain, C. & Fuchs, E. 2012. Identification of stem cell populations in sweat glands and ducts reveals roles in homeostasis and wound repair. Cell, 150, 136–50.

Luijsterburg, M. S., Dinant, C., Lans, H., Stap, J., Wiernasz, E., Lagerwerf, S., Warmerdam, D. O., Lindh, M., Brink, M. C., Dobrucki, J. W., Aten, J. A., Fousteri, M. I., Jansen, G., Dantuma, N. P., Vermeulen, W., Mullenders, L. H., Houtsmuller, A. B., Verschure, P. J. & Van Driel, R. 2009. Heterochromatin protein 1 is recruited to various types of Dna damage. J Cell Biol, 185, 577–86.

Maga, G. & Hubscher, U. 2003. Proliferating cell nuclear antigen (Pcna): a dancer with many partners. J Cell Sci, 116, 3051–60.

Mailand, N., Gibbs-Seymour, I. & Bekker-Jensen, S. 2013. Regulation of Pcna-protein interactions for genome stability. Nat Rev Mol Cell Biol, 14, 269–82.

Moldovan, G. L., Pfander, B. & Jentsch, S. 2007. Pcna, the maestro of the replication fork. Cell, 129, 665–79.

Naruse, K., Hori, H., Shimizu, N., Kohara, Y. & Takeda, H. 2004. Medaka genomics: a bridge between mutant phenotype and gene function. Mech Dev, 121, 619–28.

Nguyen, V., Deschet, K., Henrich, T., Godet, E., Joly, J. S., Wittbrodt, J., Chourrout, D. & Bourrat, F. 1999. Morphogenesis of the optic tectum in the medaka (Oryzias latipes): a morphological and molecular study, with special emphasis on cell proliferation. J Comp Neurol, 413, 385–404.

Nielsen, A. L., Oulad-Abdelghani, M., Ortiz, J. A., Remboutsika, E., Chambon, P. & Losson, R. 2001. Heterochromatin formation in mammalian cells: interaction between histones and HP1 proteins. Mol Cell, 7, 729–39.

Nowak, J. A. & Fuchs, E. 2009. Isolation and culture of epithelial stem cells. Methods Mol Biol, 482, 215–32.

Nowak, J. A., Polak, L., Pasolli, H. A. & Fuchs, E. 2008. Hair follicle stem cells are specified and function in early skin morphogenesis. Cell Stem Cell, 3, 33–43.

Oki, T., Nishimura, K., Kitaura, J., Togami, K., Maehara, A., Izawa, K., Sakaue-Sawano, A., Niida, A., Miyano, S., Aburatani, H., Kiyonari, H., Miyawaki, A. & Kitamura, T. 2014. A novel cell-cycle-indicator, mVenus-p27K-, identifies quiescent cells and visualizes G0-G1 transition. Sci Rep, 4, 4012.

Ota, S., Hisano, Y., Ikawa, Y. & Kawahara, A. 2014. Multiple genome modifications by the Crispr/Cas9 system in zebrafish. Genes Cells, 19, 555–64.

Paix, A., Folkmann, A., Goldman, D. H., Kulaga, H., Grzelak, M. J., Rasoloson, D., Paidemarry, S., Green, R., Reed, R. R. & Seydoux, G. 2017a. Precision genome editing using synthesis-dependent repair of Cas9-induced Dna breaks. Proc Natl Acad Sci U S A, 114, E10745–E10754.

Paix, A., Folkmann, A., Rasoloson, D. & Seydoux, G. 2015. High Efficiency, Homology-Directed Genome Editing in Caenorhabditis elegans Using Crispr-Cas9 Ribonucleoprotein Complexes. Genetics, 201, 47–54.

Paix, A., Folkmann, A. & Seydoux, G. 2017b. Precision genome editing using Crispr-Cas9 and linear repair templates in C. elegans. Methods, 121-122, 86–93.

Paix, A., Rasoloson, D., Folkmann, A. & Seydoux, G. 2019. Rapid Tagging of Human Proteins with Fluorescent Reporters by Genome Engineering using Double-Stranded Dna Donors. Curr Protoc Mol Biol, 129, e102.

Paix, A., Schmidt, H. & Seydoux, G. 2016. Cas9-assisted recombineering in C. elegans: genome editing using in vivo assembly of linear DNAs. Nucleic Acids Res, 44, e128.

Paix, A., Wang, Y., Smith, H. E., Lee, C. Y., Calidas, D., Lu, T., Smith, J., Schmidt, H., Krause, M. W. & Seydoux, G. 2014. Scalable and versatile genome editing using linear DNAs with microhomology to Cas9 Sites in Caenorhabditis elegans. Genetics, 198, 1347–56.

Pinto-Teixeira, F., Viader-Llargues, O., Torres-Mejia, E., Turan, M., Gonzalez-Gualda, E., Pola-Morell, L. & Lopez-Schier, H. 2015. Inexhaustible hair-cell regeneration in young and aged zebrafish. Biol Open, 4, 903–9.

Piwko, W., Olma, M. H., Held, M., Bianco, J. N., Pedrioli, P. G., Hofmann, K., Pasero, P., Gerlich, D. W. & Peter, M. 2010. RNAi-based screening identifies the Mms22L-Nfkbil2 complex as a novel regulator of Dna replication in human cells. Embo J, 29, 4210–22.

Protter, D. S. W. & Parker, R. 2016. Principles and Properties of Stress Granules. Trends Cell Biol, 26, 668–679.

Rausch, T., Hsi-Yang Fritz, M., Korbel, J. O. & Benes, V. 2019. Alfred: interactive multi-sample Bam alignment statistics, feature counting and feature annotation for long- and short-read sequencing. Bioinformatics, 35, 2489–2491.

Rausch, T., Zichner, T., Schlattl, A., Stutz, A. M., Benes, V. & Korbel, J. O. 2012. Delly: structural variant discovery by integrated paired-end and split-read analysis. Bioinformatics, 28, i333–i339.

Rembold, M., Lahiri, K., Foulkes, N. S. & Wittbrodt, J. 2006. Transgenesis in fish: efficient selection of transgenic fish by co-injection with a fluorescent reporter construct. Nat Protoc, 1, 1133–9.

Rhee, H., Polak, L. & Fuchs, E. 2006. Lhx2 maintains stem cell character in hair follicles. Science, 312, 1946–9.

Romero-Carvajal, A., Navajas Acedo, J., Jiang, L., Kozlovskaja-Gumbriene, A., Alexander, R., Li, H. & Piotrowski, T. 2015. Regeneration of Sensory Hair Cells Requires Localized Interactions between the Notch and Wnt Pathways. Dev Cell, 34, 267–82.

Sakaue-Sawano, A., Kurokawa, H., Morimura, T., Hanyu, A., Hama, H., Osawa, H., Kashiwagi, S., Fukami, K., Miyata, T., Miyoshi, H., Imamura, T., Ogawa, M., Masai, H. & Miyawaki, A. 2008. Visualizing spatiotemporal dynamics of multicellular cell-cycle progression. Cell, 132, 487–98.

Santos, A., Wernersson, R. & Jensen, L. J. 2015. Cyclebase 3.0: a multi-organism database on cell-cycle regulation and phenotypes. Nucleic Acids Res, 43, D1140–4.

Schindelin, J., Arganda-Carreras, I., Frise, E., Kaynig, V., Longair, M., Pietzsch, T., Preibisch, S., Rueden, C., Saalfeld, S., Schmid, B., Tinevez, J. Y., White, D. J., Hartenstein, V., Eliceiri, K., Tomancak, P. & Cardona, A. 2012. Fiji: an open-source platform for biological-image analysis. Nat Methods, 9, 676–82.

Seleit, A., Gross, K., Onistschenko, J., Woelk, M., Autorino, C. & Centanin, L. 2020. Development and regeneration dynamics of the Medaka notochord. Dev Biol, 463, 11–25.

Seleit, A., Kramer, I., Ambrosio, E., Dross, N., Engel, U. & Centanin, L. 2017a. Sequential organogenesis sets two parallel sensory lines in medaka. Development, 144, 687–697.

Seleit, A., Kramer, I., Riebesehl, B. F., Ambrosio, E. M., Stolper, J. S., Lischik, C. Q., Dross, N. & Centanin, L. 2017b. Neural stem cells induce the formation of their physical niche during organogenesis. Elife, 6.

Sellers, J. R. 2000. Myosins: a diverse superfamily. Biochim Biophys Acta, 1496, 3–22.

Shaner, N. C., Lambert, G. G., Chammas, A., Ni, Y., Cranfill, P. J., Baird, M. A., Sell, B. R., Allen, J. R., Day, R. N., Israelsson, M., Davidson, M. W. & Wang, J. 2013. A bright monomeric green fluorescent protein derived from Branchiostoma lanceolatum. Nat Methods, 10, 407–9.

Shin, J., Chen, J. & Solnica-Krezel, L. 2014. Efficient homologous recombination-mediated genome engineering in zebrafish using Tale nucleases. Development, 141, 3807–18.

Snippert, H. J., Van Der Flier, L. G., Sato, T., Van Es, J. H., Van Den Born, M., Kroon-Veenboer, C., Barker, N., Klein, A. M., Van Rheenen, J., Simons, B. D. & Clevers, H. 2010. Intestinal crypt homeostasis results from neutral competition between symmetrically dividing Lgr5 stem cells. Cell, 143, 134–44.

Stemmer, M., Thumberger, T., Del Sol Keyer, M., Wittbrodt, J. & Mateo, J. L. 2015. CCTop: An Intuitive, Flexible and Reliable Crispr/Cas9 Target Prediction Tool. PLoS One, 10, e0124633.

Stenmark, H. 2009. Rab GTPases as coordinators of vesicle traffic. Nat Rev Mol Cell Biol, 10, 513–25.

Stolper, J., Ambrosio, E. M., Danciu, D. P., Buono, L., Elliott, D. A., Naruse, K., Martinez-Morales, J. R., Marciniak-Czochra, A. & Centanin, L. 2019. Stem cell topography splits growth and homeostatic functions in the fish gill. Elife, 8.

Stuart, G. W., Vielkind, J. R., Mcmurray, J. V. & Westerfield, M. 1990. Stable lines of transgenic zebrafish exhibit reproducible patterns of transgene expression. Development, 109, 577–84.

Sugiyama, M., Sakaue-Sawano, A., Iimura, T., Fukami, K., Kitaguchi, T., Kawakami, K., Okamoto, H., Higashijima, S. & Miyawaki, A. 2009. Illuminating cell-cycle progression in the developing zebrafish embryo. Proc Natl Acad Sci U S A, 106, 20812–7.

Suzuki, S. C. & Takeichi, M. 2008. Cadherins in neuronal morphogenesis and function. Dev Growth Differ, 50 Suppl 1, S119–30.

Taylor, K. C., Buvoli, M., Korkmaz, E. N., Buvoli, A., Zheng, Y., Heinze, N. T., Cui, Q., Leinwand, L. A. & Rayment, I. 2015. Skip residues modulate the structural properties of the myosin rod and guide thick filament assembly. Proc Natl Acad Sci U S A, 112, E3806–15.

Thacker, S. A., Bonnette, P. C. & Duronio, R. J. 2003. The contribution of E2F-regulated transcription to Drosophila Pcna gene function. Curr Biol, 13, 53–8.

Thorvaldsdottir, H., Robinson, J. T. & Mesirov, J. P. 2013. Integrative Genomics Viewer (Igv): high-performance genomics data visualization and exploration. Brief Bioinform, 14, 178–92.

Tsingos, E., Hockendorf, B., Sutterlin, T., Kirchmaier, S., Grabe, N., Centanin, L. & Wittbrodt, J. 2019. Retinal stem cells modulate proliferative parameters to coordinate post-embryonic morphogenesis in the eye of fish. Elife, 8.

Wada, H., Dambly-Chaudiere, C., Kawakami, K. & Ghysen, A. 2013. Innervation is required for sense organ development in the lateral line system of adult zebrafish. Proc Natl Acad Sci U S A, 110, 5659–64.

Wang, H., La Russa, M. & Qi, L. S. 2016. Crispr/Cas9 in Genome Editing and Beyond. Annu Rev Biochem, 85, 227–64.

Welz, T., Wellbourne-Wood, J. & Kerkhoff, E. 2014. Orchestration of cell surface proteins by Rab11. Trends Cell Biol, 24, 407–15.

Wheeler, J. R., Matheny, T., Jain, S., Abrisch, R. & Parker, R. 2016. Distinct stages in stress granule assembly and disassembly. Elife, 5.

Wierson, W. A., Simone, B. W., Warejoncas, Z., Mann, C., Welker, J. M., Kar, B., Emch, M. J., Friedberg, I., Gendron, W. A. C., Barry, M. A., Clark, K. J., Dobbs, D. L., Mcgrail, M. A., Ekker, S. C. & Essner, J. J. 2019. Expanding the Crispr Toolbox with ErCas12a in Zebrafish and Human Cells. Crispr J, 2, 417–433.

Wierson, W. A., Welker, J. M., Almeida, M. P., Mann, C. M., Webster, D. A., Torrie, M. E., Weiss, T. J., Kambakam, S., Vollbrecht, M. K., Lan, M., Mckeighan, K. C., Levey, J., Ming, Z., Wehmeier, A., Mikelson, C. S., Haltom, J. A., Kwan, K. M., Chien, C. B., Balciunas, D., Ekker, S. C., Clark, K. J., Webber, B. R., Moriarity, B. S., Solin, S. L., Carlson, D. F., Dobbs, D. L., Mcgrail, M. & Essner, J. 2020. Efficient targeted integration directed by short homology in zebrafish and mammalian cells. Elife, 9.

Winkler, C., Vielkind, J. R. & Schartl, M. 1991. Transient expression of foreign Dna during embryonic and larval development of the medaka fish (Oryzias latipes). Mol Gen Genet, 226, 129–40.

Won, M. & Dawid, I. B. 2017. Pcr artifact in testing for homologous recombination in genomic editing in zebrafish. PLoS One, 12, e0172802.

Yamaguchi, M., Hayashi, Y., Nishimoto, Y., Hirose, F. & Matsukage, A. 1995. A nucleotide sequence essential for the function of Dre, a common promoter element for Drosophila DNa replication-related genes. J Biol Chem, 270, 15808–14.

Yan, B. W., Zhao, Y. F., Cao, W. G., Li, N. & Gou, K. M. 2013. Mechanism of random integration of foreign Dna in transgenic mice. Transgenic Res, 22, 983–92.

Yang, P., Mathieu, C., Kolaitis, R. M., Zhang, P., Messing, J., Yurtsever, U., Yang, Z., Wu, J., Li, Y., Pan, Q., Yu, J., Martin, E. W., Mittag, T., Kim, H. J. & Taylor, J. P. 2020. G3BP1 Is a Tunable Switch that Triggers Phase Separation to Assemble Stress Granules. Cell, 181, 325–345 e28.

Yao, X., Wang, X., Hu, X., Liu, Z., Liu, J., Zhou, H., Shen, X., Wei, Y., Huang, Z., Ying, W., Wang, Y., Nie, Y. H., Zhang, C. C., Li, S., Cheng, L., Wang, Q., Wu, Y., Huang, P., Sun, Q., Shi, L. & Yang, H. 2017. Homology-mediated end joining-based targeted integration using Crispr/Cas9. Cell Res, 27, 801–814.

Yoshimi, K., Kunihiro, Y., Kaneko, T., Nagahora, H., Voigt, B. & Mashimo, T. 2016. ssODN-mediated knock-in with Crispr-Cas for large genomic regions in zygotes. Nat Commun, 7, 10431.

Zerjatke, T., Gak, I. A., Kirova, D., Fuhrmann, M., Daniel, K., Gonciarz, M., Muller, D., Glauche, I. & Mansfeld, J. 2017. Quantitative Cell Cycle Analysis Based on an Endogenous All-in-One Reporter for Cell Tracking and Classification. Cell Rep, 19, 1953–1966.

Zu, Y., Tong, X., Wang, Z., Liu, D., Pan, R., Li, Z., Hu, Y., Luo, Z., Huang, P., Wu, Q., Zhu, Z., Zhang, B. & Lin, S. 2013. Talen-mediated precise genome modification by homologous recombination in zebrafish. Nat Methods, 10, 329–31.

